# Adaptation of pancreatic cancer cells to nutrient deprivation is reversible and requires glutamine synthetase stabilization by mTORC1

**DOI:** 10.1101/2020.02.16.951681

**Authors:** Pei-Yun Tsai, Min-Sik Lee, Unmesh Jadhav, Insia Naqvi, Shariq Madha, Ashley Adler, Meeta Mistry, Sergey Naumenko, Caroline A. Lewis, Daniel S. Hitchcock, Frederick R. Roberts, Peter DelNero, Thomas Hank, Kim C. Honselmann, Vicente Morales Oyarvide, Mari Mino-Kenudson, Clary B. Clish, Ramesh A. Shivdasani, Nada Y. Kalaany

## Abstract

Pancreatic ductal adenocarcinoma (PDA) is a lethal, therapy-resistant cancer that thrives in a highly desmoplastic, nutrient-deprived microenvironment. Several studies investigated the effects of depriving PDA of either glucose or glutamine alone. However, the consequences on PDA growth and metabolism of limiting both preferred nutrients have remained largely unknown. Here, we report the selection for clonal human PDA cells that survive and adapt to limiting levels of both glucose and glutamine. We find that adapted clones exhibit increased growth *in vitro* and enhanced tumor-forming capacity *in vivo*. Mechanistically, adapted clones share common transcriptional and metabolic programs, including amino acid use for *de novo* glutamine and nucleotide synthesis. They also display enhanced mTORC1 activity that prevents the proteasomal degradation of glutamine synthetase (GS), the rate-limiting enzyme for glutamine synthesis. This phenotype is notably reversible, with PDA cells acquiring alterations in open chromatin upon adaptation. Silencing of GS suppresses the enhanced growth of adapted cells and mitigates tumor growth. These findings identify non-genetic adaptations to nutrient deprivation in PDA and highlight GS as a dependency that could be targeted therapeutically in pancreatic cancer patients.

**Significance:** Pancreatic ductal adenocarcinoma (PDA) is a highly lethal malignancy with no effective therapies. PDA aggressiveness partly stems from its ability to grow within a uniquely dense stroma restricting nutrient access. This study demonstrates that PDA clones that survive chronic nutrient deprivation acquire reversible non-genetic adaptations allowing them to switch between metabolic states optimal for growth under nutrient-replete or nutrient-deprived conditions. One contributing factor to this adaptation mTORC1 activation, which stabilizes glutamine synthetase (GS) necessary for glutamine generation in nutrient-deprived cancer cells. Our findings imply that although total GS levels may not be a prognostic marker for aggressive disease, GS inhibition is of high therapeutic value, as it targets specific cell clusters adapted to nutrient starvation, thus mitigating tumor growth.

## Introduction

With a steady rise in incidence and a poor 5-year survival rate of ~ 9%, PDA is now the third leading cause of cancer death in the US (1, 2). Oncogenic *KRAS* mutations are highly prevalent in PDA (> 90%), driving many of its distinctive metabolic features (3, 4). These include the macropinocytic uptake of extracellular macromolecules, the recycling of intracellular components (5-8), and the re-direction of glycolytic and glutaminolytic intermediates into anabolic and redox-controlling pathways required for PDA growth (9, 10). However, whether these alterations are sufficient to account for the ability of PDA to thrive in a seemingly hostile *in vivo* microenvironment, where abnormal vasculature and exuberant stroma restrict nutrient access, remains unknown.

Because nutrient availability largely dictates metabolic behavior (11), the relevance of studying tumor metabolism within the native environment has been recently underscored (12-16). Although culture media composition can be modulated to mimic circulating metabolite levels (17, 18), this modulation may not accurately reflect the tumor metabolic microenvironment. In particular, levels of glucose and glutamine, two of the most abundant and tumor-preferred nutrients in the circulation, can be limiting in PDA tumors, compared to benign adjacent tissue (7), and significantly lower within tumor cores, compared to the periphery (19, 20). Paradoxically, both glucose and glutamine are routinely supplemented in culture media at levels that are significantly higher than the circulation: 11mM glucose and 2mM glutamine in RPMI compared to ~ 5.5mM glucose and 0.6mM glutamine in serum (21). Although prior studies focused on depriving tumor cells of either glucose or glutamine alone (10, 22–25), how PDA cells survive and proliferate under limiting levels of both major nutrients is not well understood.

To investigate this, we selected for clonal PDA cells that survive and adapt to limiting levels of both glucose and glutamine. We find that the adapted clones have enhanced proliferation *in vitro* and tumor-forming capacity *in vivo*. They also share common signaling, transcriptional and metabolic alterations that are acquired upon adaptation. These include a post-translational role for mTORC1 signaling in the stabilization of glutamine synthetase (GS), and the use of amino acids for the synthesis of glutamine and nucleotides. Of note, this phenotype is reversible, implicating epigenetic alterations in the adaptation process, and highlighting GS as a candidate for therapeutic targeting in pancreatic cancer.

## Results

### PDA cells adapted to low glucose / low glutamine have enhanced growth *in vitro* and *in vivo*

To test whether pancreatic cancer cells can survive and adapt to harsh nutrient-deprived conditions, we cultured seven human PDA cell lines (AsPC-1, BxPC-3, HPAC, MIA PaCa-2, PANC-1, SUIT-2 and PA-TU-8988T, herein termed 8988T) in customized low glucose (0.5mM) and low glutamine (0.1mM), medium (L-L). These modifications represent ~20-fold reduction from standard RPMI medium, herein referred to as high glucose-high glutamine (H-H) and 5- to 10-fold decrease from human serum levels (21) (Fig. 1*A*); both L-L and H-H media were supplemented with 10% dialyzed fetal bovine serum (FBS). Consistent with the requirement of glucose and glutamine for PDA growth (9, 10), most PDA cells died within two weeks of culture in L-L medium, despite media replenishment every 2-3 days (Fig. 1*B* and Fig. S1*A*). However, rare surviving cells were able to slowly proliferate past the second week, particularly in SUIT-2 and 8988T, followed by MIA-PaCa-2 (Fig. 1*B* and and Fig. S1*B*). Distinct genetic lesions frequently found in PDAC did not explain the differential adaptive capacity of these cell lines to nutrient deprivation (Fig. S1*C*). The surviving cells in SUIT-2 and 8988T gradually gave rise to adapted clones, 3 of which (A-C4, C-5 and C-6) were selected for further analysis (Fig. 1*A*). Adapted clones were then studied in comparison to non-adapted clones (NA-C1, C-2 and C-3), which were isolated from the corresponding parental PDA lines grown under H-H conditions (Fig. 1*A*). All clones were subsequently passaged every 3-4 days for use in experiments.

**Figure 1.**
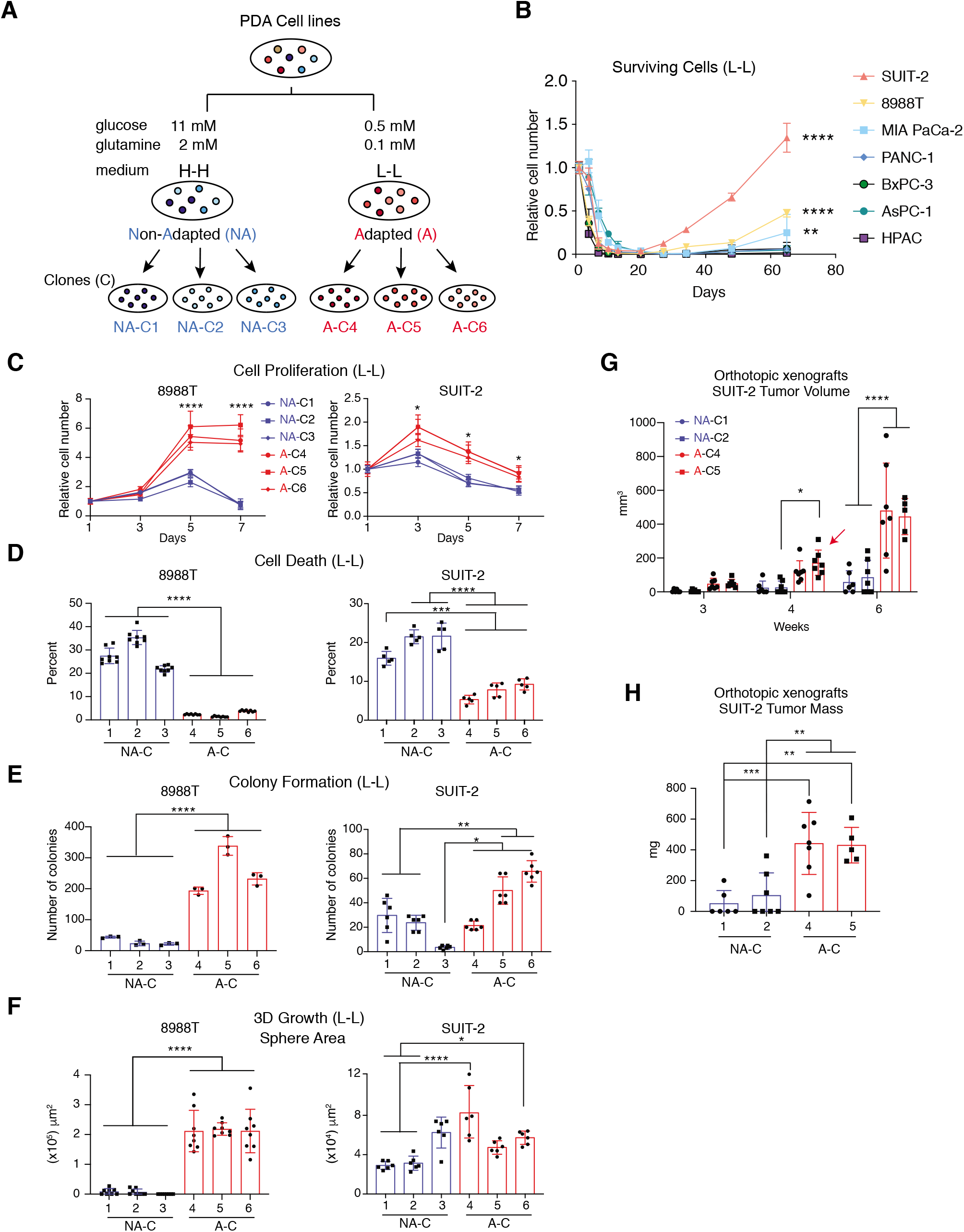
PDA cells adapted to nutrient deprivation exhibit enhanced growth *in vitro* and *in vivo*. (*A*) Schematic depicting the adaptation process to glucose and glutamine deprivation, followed by selection of adapted “A” clones (C4-C6) derived from SUIT-2 or 8988T human PDA cells that survived and grew in low glucose-low glutamine (L-L) medium, and control non-adapted “NA” clones (C1-C3) derived from parental cells plated at low dilution in high glucose-high glutamine (H-H) medium. (*B*) Relative number of surviving cells from all 7 PDA cell lines that were subjected to L-L medium for a period of 65 days (n = 3). (*C*) Proliferation curves of clonal cells described in *A* that were grown in L-L medium for 7 days (n=5 for SUIT-2 and n=8 for 8988T). (*D*) Percent cell death in clones described in *A* (n=5 for SUIT-2 and n=8 for 8988T) that was quantified on Day 4 (SUIT-2) or Day 5 (8988T) of growth in L-L medium. (*E*) Colony formation assay showing number of colonies formed by SUIT-2 or 8988T clones described in *A* that were grown in L-L medium for 7 days (n=6 for SUIT-2 and n=3 for 8988T). (*F*) Three-dimensional (3D) growth assay showing area of spheres formed by SUIT-2 or 8988T cells described in *A* that were grown for 7 days or 10 days, respectively, in L-L medium supplemented with 4% matrigel (n=6 for SUIT-2 and n=8 for 8988T). (*G*) Volumes of orthotopic xenograft PDA tumors quantified by ultrasound that were derived from SUIT-2 clones described in *A* and were injected (750 × 10^3^ cells) into the pancreas of 4-6 week old Rag1^-/-^ mice. Tumors were grown for 43 days (n=7 except for NA-C1, n=6); red arrow points at the 2 largest A-C5 tumors at week 4 from mice that died before the week 6 endpoint. (*H*) Weights of orthotopic xenograft SUIT-2 tumors described in *G* that were analyzed at the endpoint, 6 weeks post-tumor cell injection. Data represent the mean ± S.E.M. in *B* and *C*, or mean ± S.D. in *D-H*. \**p* < 0.05; \*\**p* < 0.01; \*\*\**p* < 0.001; \*\*\*\**p* < 0.0001, two-way ANOVA for *B, C* and G and one-way ANOVA for *D-F* and *H*, followed by Tukey test.

To assess their *in vitro* growth potential, adapted and non-adapted cells were first subjected to a 7-day proliferation assay performed without media replenishment. Whereas no significant differences were detected in H-H medium conditions (Fig. S2*A* and *B*), adapted cells from both lines showed enhanced proliferative capacity (Fig. 1*C*) and decreased cell death (Fig. 1*D*), compared to non-adapted clones, in L-L medium. In a colony formation assay, adapted clones tended to generate a larger number of colonies in L-L but not in H-H medium (Fig. 1*E* and Fig. S2*C*). When plated under 3-dimensional (3-D) culture conditions, adapted 8988T cells formed a significantly higher number of spheres (Fig. S2*D*) with a larger area (Fig. 1*F*) than non-adapted clones in L-L, but not H-H medium (Fig. S2*E* and *F*). SUIT-2 clones formed a similar number of spheres, independent of the medium or adaptation (Fig. S2*D* and *E*). However, SUIT-2 spheres tended to be larger in adapted, compared to non-adapted clones, when cultured under L-L, but not H-H conditions (Fig. 1*F* and Fig. S2*F*).

Because autophagy and macropinocytosis are well-recognized dependencies in PDA (3, 5, 6, 8), we assessed their potential contribution to the enhanced proliferation of the adapted clones under nutrient-limiting conditions. No differential increase in autophagic flux was observed in adapted cells, compared to non-adapted controls in L-L medium (Fig. S3*A*). Moreover, uptake of tetramethylrhodamine (TMR)-labeled dextran, an indicator of macropinocytosis, was not further increased in the adapted, compared to non-adapted cells, under L-L conditions (Fig. S3*B*). These data indicate that surviving PDA cells that adapt to chronic glucose and glutamine deprivation acquire a proliferative potential that depends on processes other than autophagy or macropinocytosis.

To test whether the adaptation-mediated *in vitro* phenotype could be translated *in vivo*, adapted and non-adapted SUIT-2 PDA clonal cells were orthotopically injected into the pancreas of B6.Rag1^-/-^ mice (Fig. 1*G* and *H* and Fig. S2*G*). Within the span of 6 weeks, only 8 out of 13 non-adapted cell injections derived from 2 different clones formed detectable, yet smaller tumors (~ 119 mm^3^ in volume; 197 mg in weight) with delayed latency (Fig. 1*G* and *H*). In contrast, 14 of 14 adapted clonal cell injections produced tumors that were detected as early as week 3 by ultrasound (Fig. 1*G*). On week 6, these tumors were approximately 4-fold larger in volume (~ 463 mm^3^) and over 2-fold heavier (~ 436 mg) than tumors from non-adapted clones (Fig. 1*G* and *H* and Fig. S1*G*). Moreover, two mice bearing the largest tumors from adapted cells (Fig. 1*G*, clone 5, red arrow) succumbed to tumor burden on week 5 and were therefore not included in the analyses at week 6 (Fig. 1*G* and *H*). These results demonstrate that metabolic adaptation to a nutrient-deprived environment can endow cancer cells with higher growth fitness, allowing tumors to survive and thrive during periods of nutrient starvation both *in vitro* and *in vivo*.

### Adapted PDA clones display altered amino acid and nucleotide metabolism coupled with enhanced mTORC1 signaling

To identify transcriptional alterations that convey enhanced proliferative potential to PDA upon adaptation to nutrient deprivation, we used RNA-Seq to profile transcripts in SUIT-2 clones under L-L conditions. Gene set enrichment analysis (GSEA) revealed significant positive enrichment in the adapted clones for pathways involved in purine and pyrimidine metabolism (Fig. S4*A* and *B*) and amino acid metabolism (Fig. S4*C-F*), and negative enrichment for genes that suppress or are negatively correlated with the mammalian target of rapamycin complex 1 (mTORC1) signaling pathway (Fig. S4*G*). Metabolic profiling of SUIT-2 clones treated for 24 hours with H-H or L-L medium yielded an adaptation signature that largely overlaps with the altered transcriptomes (Fig. S5*A* and *B*). Compared to H-H, all clones under L-L conditions displayed markedly higher levels of intracellular amino acids (Fig. S5*A*), except for alanine, glutamine, glutamate and aspartate, consistent with glutamine deprivation and indicative of enhanced autophagy. This increase in amino acids was significantly suppressed in the adapted clones, compared to non-adapted controls, suggesting enhanced utilization of intracellular amino acids (Fig. 3*E*, relative levels between 2 groups in L-L and Fig. S5*A*, relative levels among all 4 groups in L-L and H-H). Furthermore, non-adapted cells showed significant accumulation of mono- and di-nucleotides and their biosynthetic precursors, reflecting decreased DNA synthesis. In contrast, the adapted clones displayed a nucleotide biosynthesis profile that closely mirrors that of PDA clones under H-H conditions, consistent with enhanced proliferation. This profile included lower levels of mono- and di-nucleotide phosphates, used to generate tri-nucleotide phosphates, which differentially accumulate in the adapted clones (Fig. 2*A*, relative levels between 2 groups in L-L and Fig. S5*A*, relative levels among 4 groups in L-L and H-H). Indeed, although *de novo* pyrimidine synthesis was higher under H-H compared to L-L conditions, a differential increase upon adaptation was observed only under L-L conditions (Fig. 2*B*). This was evident in doubling of ^14^C incorporation into DNA, 40 hours after labeling of cells with ^14^C-aspartate (Fig. 2*B*).

**Figure 2.**
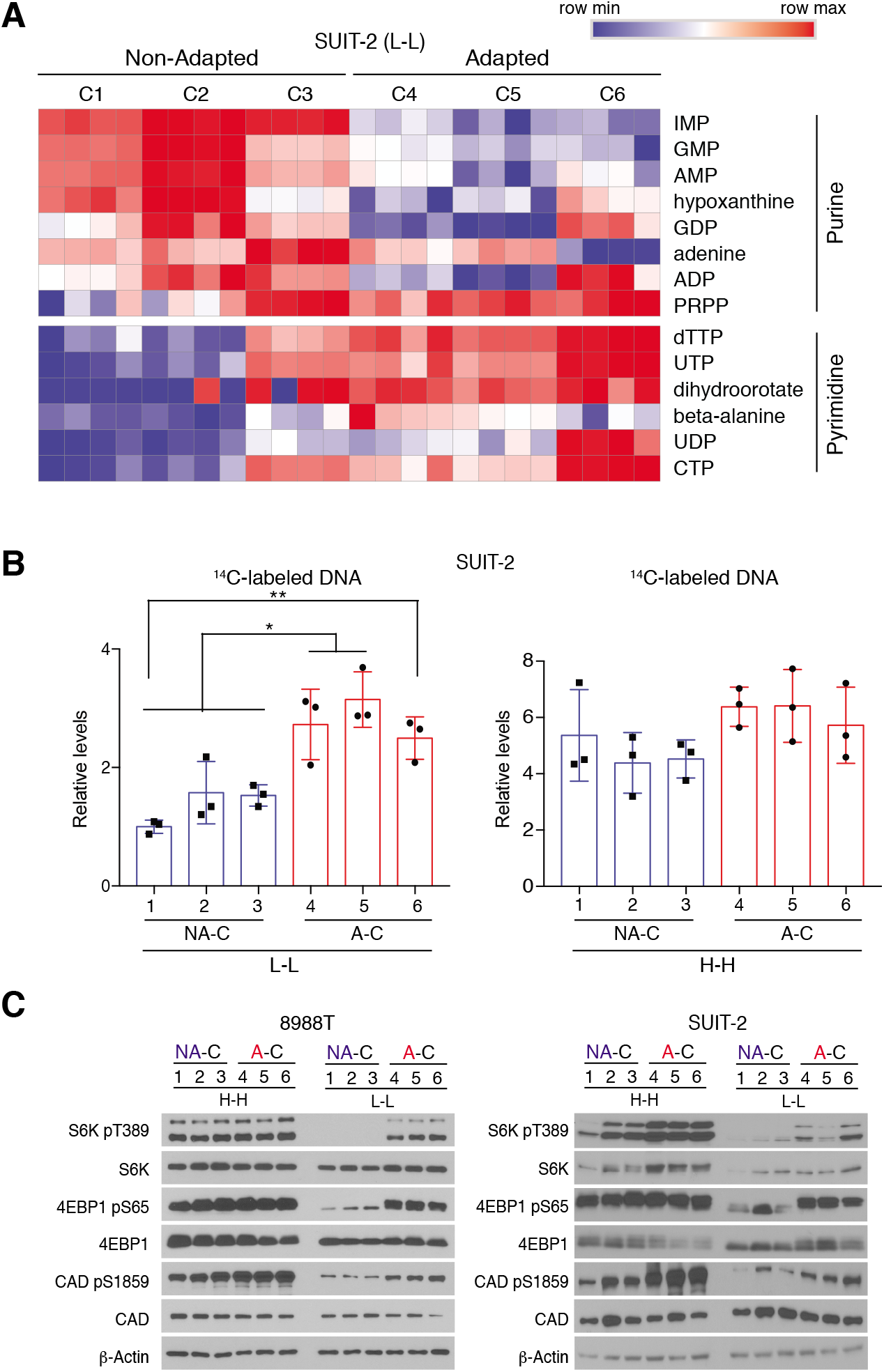
Adapted PDA clones display enhanced *de novo* DNA synthesis and mTORC1 signaling under nutrient-deprived conditions. (*A*) Heatmap listing in descending order of statistical significance (*p* < 0.05 by *t* test) metabolites in the purine and pyrimidine synthesis pathways for non-adapted (C1-C3) or adapted (C4-C5) SUIT-2 clones that were treated for 24 h with L-L medium (n=4 replicates per clone). Red indicates higher level and blue lower level, relative to the median for each metabolite across all groups. (*B*) Relative incorporation of radiolabeled U-^14^C-aspartate into DNA synthesis in non-adapted (NA-C) and adapted (A-C) SUIT-2 clones treated with either L-L or H-H medium for 40 h. Data are presented as relative fold change normalized to NA-C1 under L-L conditions and represent the mean ± S.D. (n=3 replicates per clone). \**p* < 0.05; \*\**p* < 0.01, one-way ANOVA followed by Tukey test. (*C*) Immunoblots of pT389-S6K, pS65-4EBP1, pS1859-CAD and total S6K, 4EBP1, CAD in SUIT-2 or 8988T clones treated with H-H or L-L medium for 24 h. β-actin was used as loading control.

**Figure 3.**
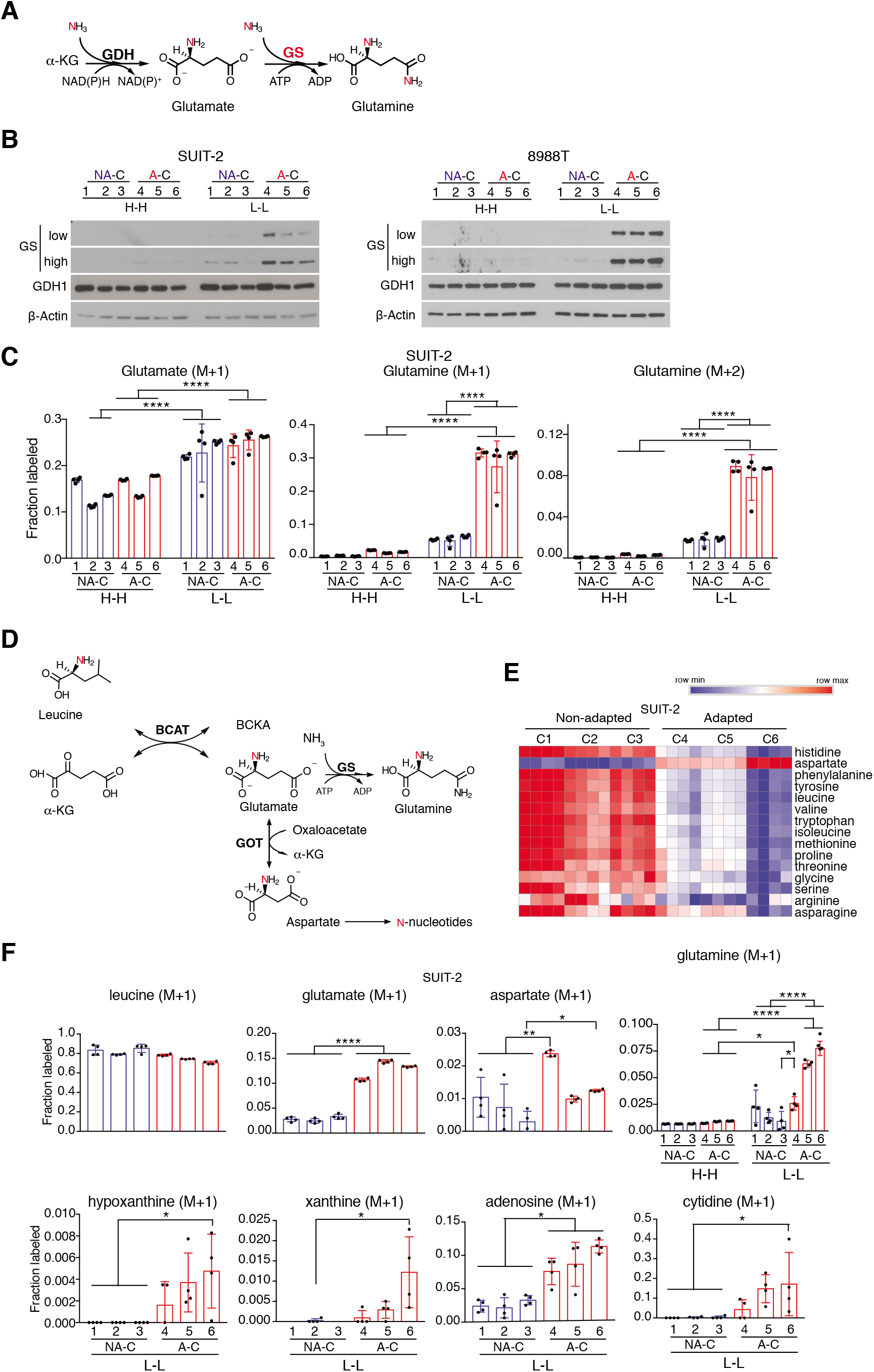
Adapted PDA cells induce GS and have enhanced incorporation of nitrogen from amino acids into glutamine and nucleotide synthesis. (*A*) Schematic illustrating glutamate synthesis from α-ketoglutarate (α-KG) and ammonia by glutamate dehydrogenase (GDH) and glutamine synthesis from glutamate and ammonia by glutamine synthetase (GS). In red, are ammonia nitrogens incorporating into glutamate and then glutamine. (*B*) Immunoblots of GS (low and high exposures) and GDH in SUIT-2 or 8988T clones treated with H-H or L-L medium for 24 h. β-actin was used as loading control. (*C*) Fractional labeling of ^15^N-glutamate and glutamine in SUIT-2 cells treated for 24 h with H-H or L-L medium supplemented with 0.8mM ^15^N-ammonium chloride. Data are corrected for natural abundance and represent the average of 4 replicates per clone per condition ± S.D. (*D*) Schematic illustrating the incorporation of leucine nitrogen (red) into glutamate and then aspartate via transamination reactions catalyzed by branched chain amino acid transaminase BCAT and aspartate transaminase GOT. Glutamate is used to synthesize glutamine via GS and aspartate nitrogen incorporates into newly synthesized nucleotides. (*E*) Heatmap listing in descending order of statistical significance (*p* < 0.05 by *t* test) amino acids in non-adapted or adapted SUIT-2 clones that were treated for 24 h with L-L medium (n=4 per clone per condition). Red indicates higher level and blue lower level, relative to the median for each metabolite across all groups. (*F*) Fractional labeling of metabolites in glutamine and nucleotide synthesis pathways in cells described in *B* grown in media supplemented with 0.4mM ^15^N-leucine for 24 h. Data represent the average of 4 replicates per clone per condition ± S.D. \**p* < 0.05; \*\**p* < 0.01; \*\*\*\**p* < 0.0001, two-way ANOVA for *C*, one-way ANOVA for *F* except for “Glutamine” (two-way ANOVA), followed by Tukey test.

Signaling downstream of mTORC1 can induce *de novo* pyrimidine synthesis through activation of the rate-limiting enzyme “CAD” or carbamoyl-phosphate synthetase 2, aspartate transcarbamoylase, dihydroorotase (26). Since GSEA also indicated a potential increase in mTORC1 activity in adapted clones under L-L conditions (Fig. S4*G*), we investigated a role for mTORC1 in the adaptation of the PDA clones to nutrient deprivation. Compared to non-adapted clones, adapted SUIT-2 and 8988T cells showed increased phosphorylation levels of CAD on Serine 1859, a site of ribosomal S6-Kinase (S6K1) activity (Fig. 2*C*). This increase was particularly striking under L-L conditions. Importantly, it was mirrored by enhanced mTORC1 activity, as evidenced by sustained phosphorylation of its downstream effectors S6K1 and 4E-binding protein (4E-BP1), despite nutrient deprivation (Fig. 2*C*). Consistently, the adapted clones displayed enhanced sensitivity to higher levels of the mTORC1 inhibitor rapamycin (1 μM-20μM), as demonstrated by decreased live cell number over a 48-hour treatment (Fig. S6*B*). On the other hand, no significant changes were detected in the phosphorylation levels of ERK1/2 downstream of activated KRAS. Moreover, levels of AKT phosphorylation (T308) were decreased in adapted clones (Fig. S6*A*), consistent with mTORC1-AKT negative feedback loops (27-29).

### Adapted PDA clones have enhanced glutamine synthesis

Glutamine is a pleiotropic amino acid required for protein synthesis and numerous anabolic reactions in proliferating cells (22). We therefore asked whether the adapted clones have induced *de novo* glutamine synthesis under L-L conditions. mRNA levels of glutamine synthetase (GS, also termed GLUL), which catalyzes the condensation of glutamate and ammonium to generate glutamine (Fig. 3*A*), were only moderately increased in adapted clones (Fig. S7*A*). However, GS protein levels were markedly induced in adapted clones from both PDA cell lines, compared to non-adapted controls (Fig. 3*B*). This occurred only under L-L conditions, as no clones harbored detectable levels of GS in H-H medium, independent of adaptation (Fig. 3*B*). Although these findings are consistent with a glutamine-dependent post-translational regulation of GS (30, 31), it is noteworthy that non-adapted clones only displayed a mild to no increase in GS levels 24 hours following exposure to L-L medium (Fig. 3*B*), implying differential metabolic rewiring in the PDA clones upon adaptation to prolonged glutamine deprivation. Importantly, GS protein levels were mirrored by its enzymatic activity levels, as demonstrated by metabolic tracing of ^15^N-ammonium chloride (NH_4_Cl) in SUIT-2 cells (Fig. 3*C*). Because glutamate dehydrogenase (GDH, also termed GLUD) catalyzes the reductive amination of alpha-ketoglutarate (α-KG) to generate glutamate (Fig. 3*A*), a moderate increase was observed in the ^15^N-glutamate fraction between H-H and L-L conditions in SUIT-2 cells (~18% vs 25%), independent of adaptation (Fig. 3*C*). This suggested the use by the adapted cells of newly generated glutamate to synthesize glutamine. Indeed, adapted clones cultured in L-L medium had a marked increase in ^15^N-glutamine (M+1) fraction, generated from either condensation of ^15^N-ammonium and unlabeled glutamate, or unlabeled ammonium and GDH-derived ^15^N-glutamate, compared to non-adapted controls (30% vs 8%, respectively). This increase was also observed in the (M+2) labeled fraction, which is generated when both substrates are labeled (8% in adapted vs 2% in non-adapted cells). On the other hand, ^15^N-glutamine levels were either undetectable or minimally detected in the PDA clones under H-H conditions, independent of adaptation (Fig. 3*C*). These data indicate that only PDA clones that survive and adapt to long-term glutamine deprivation are able to induce GS protein levels under L-L conditions, so as to increase GS-mediated glutamine synthesis.

### Adapted PDA cells use amino acids as a nitrogen source for glutamine and nucleotide synthesis

In addition to reductive amination of α-KG through GDH (Fig. 3*A*), the transamination reaction catalyzed by branched chain amino acid (BCAA) transaminase (BCAT1/2) could contribute to the generation of mitochondrial glutamate that is then used for glutamine synthesis. Moreover, aspartate transaminase (GOT1/2, also termed AST1/2) could transfer the nitrogen from glutamate to aspartate, a key substrate in nucleotide biosynthesis (Fig. 3*D*). Global metabolite profiling of the PDA clones revealed a significant decrease in amino acid levels, including the branched chain amino acids (BCAA) leucine, valine and isoleucine in all adapted clones, suggesting enhanced amino acid utilization by the more proliferative cells (Fig. *3E* and Fig. S5*A*). Indeed, adapted clones showed increased protein levels of BCAT2 and GOT2 in SUIT-2 cells, whereas BCAT1 and GOT1 were increased in 8988T cells (Fig. S7*B*). ^15^N-leucine tracing demonstrated that the labeled nitrogen incorporates at significantly higher levels into glutamate, aspartate and intermediates in nucleotide synthesis in adapted, compared to non-adapted clones under L-L conditions (Fig. 3*F*). Moreover, the ^15^N-labeled fraction of glutamine was minimal (1%) under nutrient-replete H-H conditions, whereas a marked 5-fold increase in ^15^N-glutamine was detected under L-L conditions, specifically in the adapted clones (Fig. 3*F*). These data imply that BCAAs are a significant source for *de novo* synthesis of glutamine and nucleotides in PDA cells adapted to L-L conditions.

### Deprivation of glutamine rather than glucose is a larger contributor to the enhanced proliferative fitness of adapted PDA cells

To distinguish the contributions of glucose versus glutamine to the adaptability phenotype, we selected for SUIT-2 and 8988T parental cells that survive and adapt to deprivation of either glucose-only (low glucose-high glutamine; L-H medium), glutamine-only (high glucose-low glutamine; H-L medium), or both glucose and glutamine (L-L). We then compared their proliferative capacity to that of control non-adapted parental cells that were maintained in nutrient replete (H-H) medium. Under H-H medium, enhanced proliferation of adapted cells was only observed on the last day of the assay (day 7), when nutrients become more depleted in the non-replenished medium (Fig. S8*A*, top panels). When grown in L-L medium however, the markedly enhanced proliferative capacity of cells adapted to L-L conditions could be mostly attributed to adaptation to glutamine deprivation, rather than deprivation of glucose. This is evident in the growth curves of H-L adapted cells that more closely mimic those of L-L adapted cells; in contrast, L-H adapted cells only displayed minimal to moderately enhanced proliferative fitness (Fig. S8*A*, bottom panels). Consistently, GS protein levels were most induced in H-L compared to L-H adapted clones in both cell lines (Fig. S8*B*).

We next assessed the effects of short-term deprivation of either glucose or glutamine, on altered metabolism, mTORC1 activity and GS levels in the L-L adapted SUIT-2 clones (Fig. 1*A*). Except for metabolites in glycolysis (marked with a star), deprivation of glutamine alone (H-L), rather than glucose (L-H), more closely mimicked the metabolic alterations observed upon deprivation of both nutrients (L-L). These include changes in levels of amino acids, TCA cycle metabolites and nucleotides (Fig. S9) Moreover, whereas loss of either glucose or glutamine or both for 24 hours, strongly suppressed mTORC1 signaling in non-adapted clones, adapted cells maintained significant mTORC1 activity even in the absence of glucose, but particularly under low glutamine conditions (H-L or L-L). Similarly, GS induction was highly dependent on both adaptation status and glutamine levels, with adapted clones displaying highest GS protein levels under H-L or L-L conditions (Fig. S8*C*, with levels normalized and quantified in *D*). This led us to investigate a possible genetic basis for induced mTORC1 activity in the adapted clones and a potential role for mTORC1 in inducing GS levels under low glutamine conditions.

### mTORC1 activation and enhanced proliferative potential are reversible in adapted PDA clones

Because the adapted clones sustain active mTORC1 signaling independent of nutrient deprivation, we asked whether the adaptation process had selected for cells with inherent mutations that constitutively activate mTORC1. We first cultured the SUIT-2 adapted clones in nutrient-replete H-H medium for 4 or more consecutive passages (a process we termed “adaptation reversal”) followed by treatment for 24 hours with L-L medium (Fig. 4*A*). Reverse-adapted (RA) clones displayed gradual suppression of mTORC1 activity (Fig. 4*B*), indicating that the enhanced anabolic signaling acquired upon adaptation is an unlikely result of genetic mutations selected for in the adapted clones. This was later confirmed with Whole Exome Sequencing (WES) analysis that found no High or Moderate impact mutations (see Methods) in any of the genes in the mTORC1 signaling pathway (Dataset S1) that was identified as enriched at the transcriptional level, by GSEA (Fig. S4*G*). Indeed, of all 23 genes (Fig. S4*G*), only 10 genetic variants exist that were however found in all clones, independent of adaptation (C1-C9, Dataset S1). Consistently, adapted and RA clones clustered according to clone number (Fig. S10*A*) rather than according to adaptation, with no High impact mutations found in whole exome variants, implying that genetic differences acquired upon reverse adaptation were minimal (Fig. S10*A* and Dataset S2).

**Figure 4.**
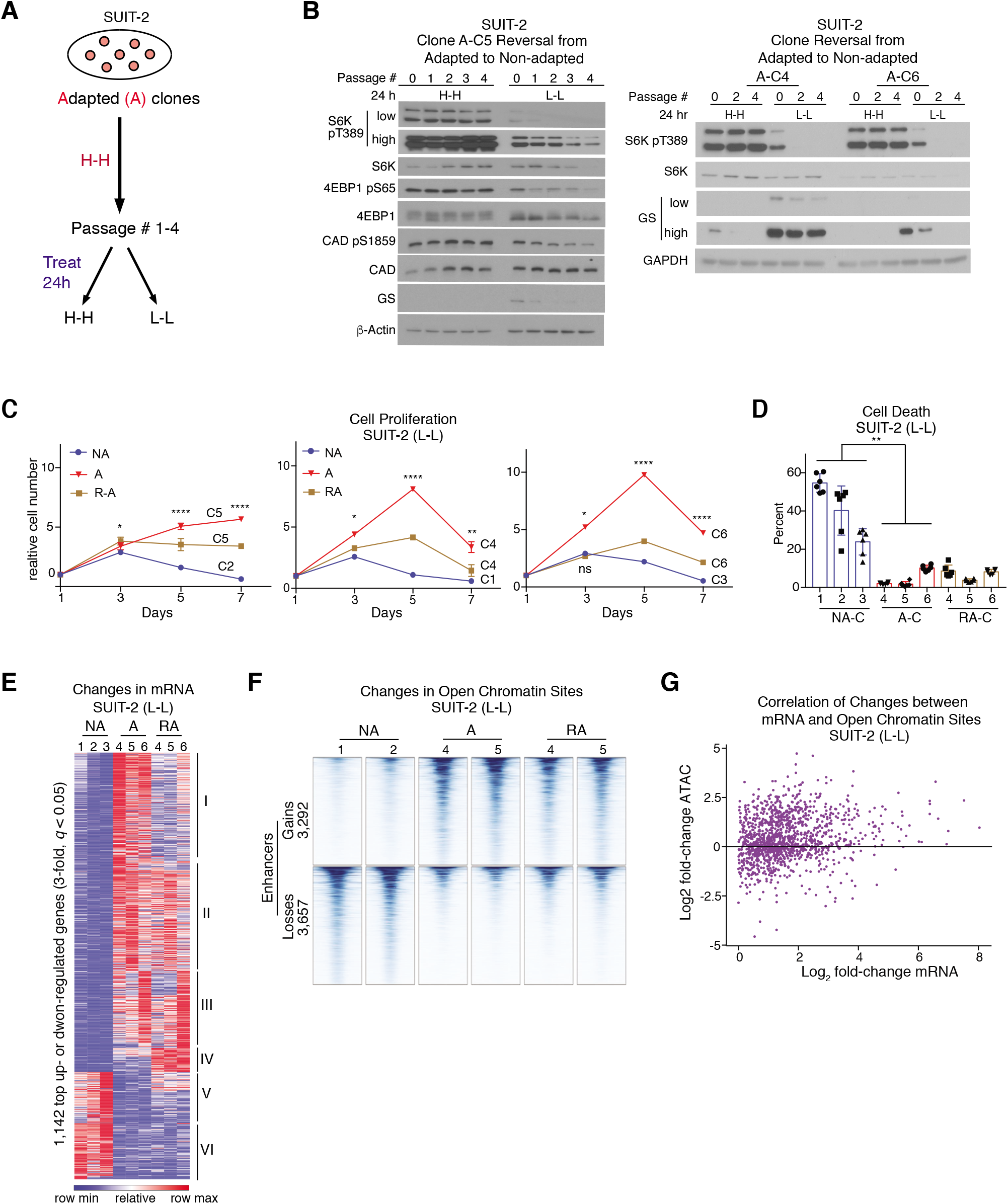
PDA adaptation to nutrient deprivation is reversible and associated with changes in open chromatin. (*A*) Schematic depicting the adaptation reversal process where adapted PDA SUIT-2 clones are grown in nutrient replete (H-H) conditions for at least 4 passages, prior to treatment with H-H or L-L medium for 24 h. (*B*) Immunoblots of pT389-S6K (low and high exposures), pS65-4EBP1, pS1859-CAD and total S6K, 4EBP1, CAD and GS (low and high exposures) in SUIT-2 clones (C5, C4 and C6) that were reverse-adapted for over 4 passages and then treated with H-H or L-L medium for 24 h. β-actin or GAPDH was used as loading control. (*C*) Proliferation curves of SUIT-2 clones C5, C4 and C6 from *B* that are either adapted “A” or reverse adapted “RA” for 18 passages, as compared to non-adapted “NA” clones C2, C1 or C3. All clones were grown under L-L conditions for 7 days without media replenishment (n=6). (*D*) Percent cell death in clones described in *C* that were grown under L-L medium for 5 days (n=6). Data represent the mean ± S.E.M. in *C*, and mean ± S.D. in *D*. \**p* < 0.05; \*\**p* < 0.01; \*\*\*\**p* < 0.0001, two-way ANOVA for *C* and one-way ANOVA for *D*, followed by Tukey test. In *C*, stars indicate statistical significance for A vs NA; A vs RA; RA vs NA on the indicated days. (*E*) Relative mRNA levels of 1,842 genes that were either upregulated or downregulated (>3-fold, *q* <0.05) in SUIT-2 adapted clones, compared to non-adapted clones; mRNA levels of the corresponding genes in reverse-adapted “RA” clones are shown alongside. Differentially expressed genes were identified by unsupervised k-means clustering. Red, high expression; blue, reduced expression relative to mean expression levels for each gene across all groups (n=3 clones per group). (*F*) ATAC-seq data showing significant (>2-fold, *q* <0.05) changes in chromatin access at enhancers (located >500 bp from transcription start sites) in each of 2 independent non-adapted (NA), adapted (A), and reverse-adapted (RA) SUIT-2 clones. (*G*) Correlation of changes in gene expression and nearby (<25 kb) chromatin accessibility for the genes that were induced upon adaptation to nutrient deprivation in SUIT-2 cells. Each dot represents a gene. More gene-linked enhancers (<25 kb) show gains than show losses in accessibility.

Importantly, *in vitro* survival and growth of adapted clones were partially reversed, with all RA clones exhibiting decreased proliferation compared to adapted clones, and one RA clone displaying a trend in enhanced cell death (Fig. 4*C* and *D*). Furthermore, among a pool of 1,842 genes whose expression was significantly altered (≥ 3-fold change, *q* <0.05) upon adaptation, about half of the changes were partially reversed in RA clones (clusters I, II and V, Fig. 4*E*), consistent with decreased mTORC1 signaling and proliferative capacity (Fig. 4*B* and *C*). These results imply that adaptation to nutrient deprivation depends on a fraction of the transcriptional changes that accompany it.

Inactive gene regulatory elements tend to be inaccessible (closed chromatin), in contrast to active cis-elements, which reside in open chromatin (32). To investigate a potential epigenetic basis for PDA cells’ adaptation to nutrient deprivation, we asked whether the transcriptional changes were associated with chromatin dynamics. Assays for transposase-accessible chromatin with high throughput sequencing (ATAC-seq) identified a significantly altered landscape of open chromatin in adapted clones, compared to non-adapted clones, including gains and losses of chromatin accessibility near thousands of enhancers (Fig. 4*F*). Adaptation-related increases in chromatin access correlated with increased mRNA levels and conversely, reduced chromatin access correlated with lower mRNA levels (Fig. 4*G* and Fig. S10*B*). Notably, alterations in open chromatin were largely preserved in reverse-adapted cells (Fig. 4*F* and Fig. S10*C*, left) with infrequent reversion (Fig. 4*F* and Fig. S10*C*, right). Thus, most changes in chromatin accessibility are stable, and again, adaptation depends only on a fraction of them. Further reversal of open chromatin changes, with corresponding changes in gene expression and near complete reversal of proliferative fitness, may require prolonged de-adaptation of PDA clones in nutrient-replete media.

### mTORC1 prevents proteasomal degradation of GS in adapted PDA clones

Because the induction of GS protein in adapted cells consistently mirrored mTORC1 signaling (Fig. 2*C*, 3*B* and Fig. S8*B* and *C*), including reversibility upon de-adaptation (Fig. 4*B*), we asked whether mTORC1 regulates GS levels. Upon treatment with the mTOR kinase inhibitor Torin1, *GS* mRNA levels were increased in adapted clones under L-L conditions (Fig. S11*A*). In contrast, consistent with post-transcriptional GS regulation, Torin1 markedly decreased GS protein levels (Fig. 5*A*). This decrease was not observed upon treatment with cycloheximide, but only when Torin1 was added to the cells, indicating that mTORC1 regulation of GS is post-translational (Fig. 5*B* and Fig. S11*B*). Indeed, whereas treatment with the proteasomal inhibitor MG-132 did not significantly affect GS levels in adapted clones over 2-8 hours, it induced GS at the 8-hour timepoint in non-adapted cells treated with L-L media (Fig. 5*C*), despite suppressed activity of the translational regulator mTORC1 under these conditions (Fig. 2*C*). Importantly, the Torin1-induced decrease in GS levels in adapted cells was rescued upon treatment with MG-132 (Fig. 5*D*). Furthermore, mTORC1 inhibition promoted GS poly-ubiquitination, as evidenced in immunoprecipitated GS from adapted cells co-treated with Torin1 and MG-132 (Fig. 5*E*). Altogether, these data demonstrate that GS is ubiquitinated and targeted for proteasomal degradation in non-adapted clones, even under L-L conditions, and that mTORC1 prevents this degradation, leading to GS stabilization in adapted clones.

**Figure 5.**
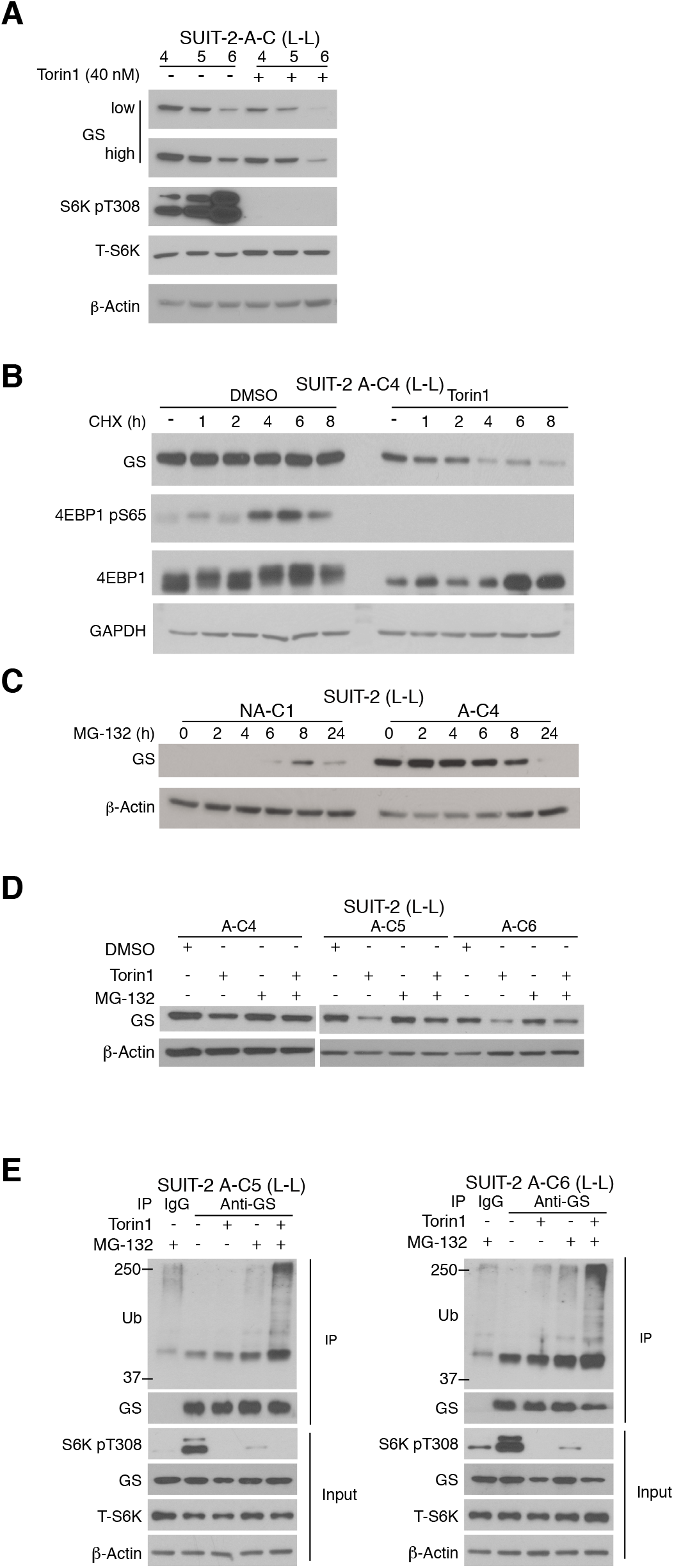
mTORC1 stabilizes GS protein levels under glutamine deprivation. (*A*) Immunoblots of GS (low and high exposures), pT389-S6K and total S6K in SUIT-2 adapted clones (C4-C6) that were treated with vehicle control (DMSO) or Torin 1 (40 nM) in L-L medium for 24 h. (*B*) Immunoblots of GS, pS65-4EBP1 and total 4EBP1 in SUIT-2 adapted clonal cells (A-C4) that were treated under L-L medium with control DMSO or Torin 1 (200 nM) for a total of 8 h, in the absence or presence of cycloheximide (CHX, 20 μg ml^-1^) for the indicated times. (*C*) GS protein levels in non-adapted (C1) and adapted (C4) SUIT-2 clones treated in L-L medium with MG-132 (10μM) for the indicated times. (*D*) GS protein levels in SUIT-2 adapted clones treated in L-L medium with either Torin 1 (200 nM) or MG-132 (10μM) alone, or in combination for 12 h (C4) or 8 h (C5, C6). (*E*) Co-immunoprecipitation (IP) blots showing the ubiquitination status of GS in SUIT-2 adapted clones treated in L-L medium, with either Torin 1 (200 nM) or MG-132 (10μM) alone, or in combination for 8 h. Input refers to immunoblots of total GS, pT389-S6K and total S6K levels in whole cell lysates. In *A-E*, β-actin or GAPDH was used as loading control.

### PDA cells adapted to nutrient-deprived conditions are sensitive to GS inhibition

To assess the contribution of glutamine synthesis to the enhanced growth of the adapted cells, PDA clones were treated with the GS inhibitor L-methionine sulfoximine (MSO). Adapted cell proliferation was mitigated under L-L, but not H-H conditions, underscoring the relevance of glutamine synthesis in the nutrient-deprived state (Fig. S12*A*). Genetic silencing of GS similarly suppressed adapted cell proliferation and colony formation (Fig. 6*A-C*, Fig. S12*B-D*). To ask whether the contribution of GS to enhanced *in vitro* PDA growth extends *in vivo*, we induced knockdown of GS in orthotopic PDA transplants (Fig. 6*D* and *E*). Five days following injection of adapted SUIT-2 cells expressing control or *GS* hairpins into the pancreas, gene silencing was induced by treating mice with doxycycline (Dox) in the drinking water. Two distinct *GS* hairpins attenuated PDA growth (Fig. 6*D* and Fig. S12*E* and *F*), resulting in 6.1- to 9.2-fold lower tumor weights upon Dox treatment (Fig. 6*D*). Although no significant changes in apoptosis were detected at the experimental endpoint, Ki-67 proliferative index tended to be moderately lower in tumors upon GS knockdown (Fig. S12*G* and *H*). Altogether, our results indicate that GS-mediated glutamine synthesis is required for PDA cells to survive and adapt to the characteristic nutrient-deprived PDA environment and highlight GS as a metabolic target in treating pancreatic cancer (Fig. 6*F*).

**Figure 6.**
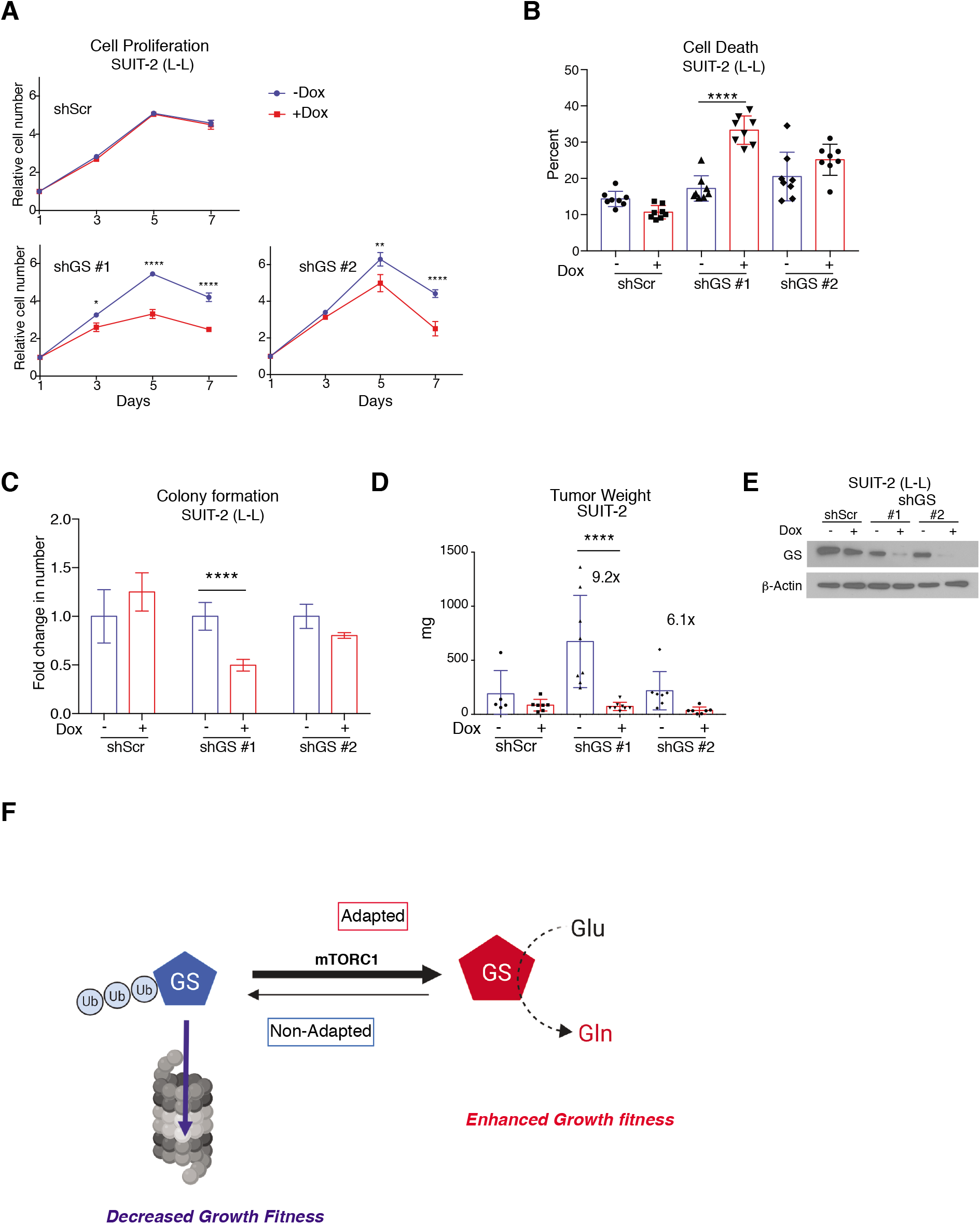
GS is required for the adaptation-induced growth fitness of PDA cells under nutrient deprivation. (*A*) Proliferation curves of adapted SUIT-2 clonal cells transfected with Doxycycline (Dox)-inducible shScrambled control or shGS hairpins 1 and 2, grown in L-L medium for 7 days in the absence or presence of 1 μg ml^-1^ Dox (n=8). (*B*) Percent cell death in cells described in *A* (n=8) that were grown in L-L medium for 7 days. (*C*) Colony formation assay for cells in *A* that were grown for 10 days, under L-L conditions (n=6). (*D*) Weights of orthotopic xenograft PDA tumors derived from SUIT-2 adapted cells (C4) stably expressing Dox-inducible hairpins described in *A* that were injected (750 × 10^3^ cells) into the pancreas of 4-6 week old Rag1^-/-^ mice. Dox was supplemented in the drinking water (2 mg ml^-1^) 5 days post-injection of cells, and tumors were harvested 33 days later (n=5 shScr-Dox; n=7 shScr + Dox; n=8 shGS#1; n=7 shGS#2). Data represent the mean ± S.E.M. in *A* and mean ± S.D. in *B-D*. \**p* < 0.05; \*\**p* < 0.01; \*\*\*\**p* < 0.0001, two-way ANOVA followed by Tukey test. (*E*) Immunoblots of GS in cells used for xenografts in *D* showing decreased protein levels, 48 h following treatment with Dox (1 μg ml^-1^) for inducible knockdown. β-actin was used as loading control. (*F*) Model illustrating a role for GS stabilization in PDA cell adaptation to nutrient deprivation, which leads to enhanced growth fitness. The sketch was created using Biorender.com.

To extend our findings to human pathology, tissue microarrays (TMAs) of resected tumors from 127 pancreatic cancer patients were immunostained and scored for GS protein levels. As a reference for intensity, stained control cores of normal pancreas showed high GS levels in islets and modest levels in acinar cells. Within tumor cores, PDA cells displayed heterogeneous cytoplasmic staining that varied widely from strong to moderate to weak, with an overall score lower than that of acinar cells. As a result, no significant association was found between PDA GS levels and tumor size, lymph node involvement (pT or pN staging, the American Joint Commission on Cancer, AJCC 8^th^ edition (33)) or patient survival (Fig. S13*A-C*). Although these results seem discordant with a recent report that GS expression is a feature of *KRAS-driven* PDA (34), the pathological heterogeneity and variability in GS protein levels is consistent with an inverse correlation with nutrient deprivation (Figs. 3, 4 and Fig. S8). Indeed, nutrient distribution is recognized to be heterogeneous in the tumor microenvironment, with glutamine levels being most depleted within tumor cores, compared to the periphery (19, 20). Therefore, future studies could confirm whether gradients of GS protein levels exist among distinct PDA cell clusters within the tumor mass, depending on their precise location and proximity to blood vessels. Nevertheless, based on our results and those of Bott *et al*. (34), targeting GS in the nutrient-deprived PDA tumors holds significant therapeutic value in pancreatic cancer.

## Discussion

The *in vitro* dependence of proliferating cells on glutamine (22) was recently questioned in the *in vivo* setting, where tumors preferably utilize glucose and other metabolites, rather than circulating glutamine, as sources of tricarboxylic acid (TCA) cycle carbon (14, 35-37). This observation is supported by *in vivo* resistance of tumors to inhibitors of glutaminolysis (38). Differences in nutrient levels between cell culture media and human serum, in addition to the *in vivo* tumor environment, might partially explain this discrepancy (12, 39). However, although glutamate is a major source of TCA cycle carbon and nitrogen for transamination reactions and glutathione synthesis, glutamine *per se* is required for protein synthesis and is an indispensable source of amide nitrogen for nucleotide and hexosamine synthesis (34). Therefore, whereas proliferating tumor cells may resort to alternative sources of glutamate, such as transamination reactions instead of glutaminolysis, they can only survive in glutamine-deprived conditions if they can synthesize glutamine *de novo*.

Moreover, although glutamine may not seem to be depleted in extracts of whole tumors or tumor interstitial fluid (16), tumor cores hold lower levels than the periphery (19, 20). Thus, the precise location of a cancer cell within the hypoxic, stroma-dense tumor may dictate its dependence on glutamine at various times. Distinct clusters of tumor cells likely exist within PDA tumors with differential nutrient access depending on the proximity to blood vessels and density of the surrounding stroma. The deprivation of PDA tumor cells of nutrients may also change over time, as tumor growth reshapes 3D cellular organization and the proximity of cancer cells to stroma and endothelial cells. It is therefore not surprising that, unlike Bott *et al*. (34), we found no significant association between average GS levels and tumor grade or patient survival (Fig. S13). We find instead that rare clonal cells are able to survive and adapt to low levels of the tumor-preferred nutrients glucose and glutamine; these cells induce glutamine synthesis, which is required for their growth *in vitro* and *in vivo*.

Despite chronic nutrient deprivation in the adapted cells, we find that activated mTORC1 induces GS. Although GS is known to be regulated post-translationally (30, 31), the signaling basis for this regulation was unknown. We show that mTORC1 plays a key post-translational role in stabilizing GS by preventing its ubiquitination and proteasomal degradation. mTORC1 inhibition was previously shown to enhance proteasomal degradation of long-lived proteins (40), but GS was not identified as a specific target, perhaps because the experiments were performed under glutamine-replete conditions. What induces mTORC1 activity in the absence of glucose and glutamine in adapted PDA cells is a question worthy of future investigation. One plausible explanation is that epigenetic alterations that occur upon adaptation to nutrient deprivation (Fig. 4 and Fig. S10) may influence genes that directly or indirectly activate mTORC1 (Fig. S4*G*).

We propose that PDA cells that survive and adapt to nutrient deprivation acquire an epigenetic state that endows them with the flexibility to switch between metabolic states that are optimal for growth under nutrient-replete or nutrient-deprived conditions. A key contributing factor to the adaptation to nutrient-depleted conditions is activation of mTORC1, which in turn stabilizes GS, thus enabling essential glutamine synthesis for pancreatic cancer growth (Fig. 6*F*). Consequently, targeting of GS, which expression is induced in nutrient-deprived cell clusters within PDA tumors, holds therapeutic benefit to pancreatic cancer patients.

## Methods

### Reagents

Antibodies for western blotting were from Abcam: BCAT1 (ab107191), GOT2 / FABP-1 (ab171739); BD Biosciences: GS / GLUL (BD-61057); Cell Signaling Technology (CST): AKT (pan, C67E7, # 4691), pS473-AKT (193H12, # 4058), pT308-AKT (244F9, # 4056), BCAT2 (# 9432), CAD (# 11933), pS1859-CAD (# 12662), 4EBP1 (53H11, # 9644), pS65-4EBP1 (# 9451), LC3B (# 2775), pThr389-S6K (# 9234); Novus Biologicals: GOT1 (NBP1-54778); Proteintech: GLUD1 / GDH1 (14299-1-AP); Santa Cruz Biotechnology: S6Kα (H-9, sc-8418), GAPDH (FL-335, sc-25778), β-Actin (sc-47778). All western blot antibodies were used at 1:1,000 dilution, except for β-Actin (1: 20,000). Antibodies for immunoprecipitation, ubiquitination or immunohistochemistry are described in the respective Methods sections below. Chloroquine (Sigma-Aldrich, C6628); Cycloheximide (Sigma-Aldrich, C4859); DMSO (Sigma-Aldrich D2650); MG-132 (Selleck Chemicals, S2619); Doxycycline (Sigma-Aldrich; D9891); L-Methionine Sulfoximine, MSO (Sigma-Aldrich; M5379); Torin 1 (Selleck Chemicals, S2827).

### Cell culture

Human PDA cell lines were from the American Type Culture Collection or ATCC (AsPC-1, BxPC-3, HPAC, MIA PaCa-2, PANC-1), Japanese Collection of Research Bioresources (SUIT-2), or the German Collection of Microorganisms and Cell Cultures (PA-TU-8988T) and were authenticated by STR profiling at ATCC. All cell lines tested negative for mycoplasma using LookOut Mycoplasma PCR Kit (Sigma, MP0035). All cells were maintained at 37°C in a humidified incubator with 5% CO2 and were grown in media supplemented with 10% dialyzed fetal bovine serum (Sigma-Aldrich F0392). For adaptation to “low glucose, low glutamine” (L-L), “low glucose-high glutamine” (L-H), or “high glucose-low glutamine” (H-L), PDA cells were cultured in RPMI 1640 medium lacking both glucose and L-glutamine (US Biological, R9011-01) that was supplemented with either 0.5 mM glucose and 0.1 mM glutamine for L-L; or with 0.5 mM glucose and 2mM glutamine (L-H); or with 11mM glucose, 0.1mM glutamine (H-L). Non-adapted cells were cultured in the same RPMI 1640 medium that was however supplemented with 11mM glucose and 2 mM glutamine, and was termed “high glucose, high glutamine” (H-H) medium. Glucose (A2494001) and glutamine (25030-81) were from Thermo Fisher Scientific.

### Gene knockdown

For small interfering RNA knockdown, ON-TARGETplus Human GS (GLUL) siRNA with target sequence: GCACACCUGUAAACGGAUA (J-008228-09) or control non-targeting siRNA: UGGUUUACAUGUUGUGUGA (D-001810-02-05) were purchased from Dharmacon, Horizon Discovery (Cambridge, UK) and transfected according to the manufacturer’s protocol. For generation of PDA cells with stable doxycycline-inducible GS or control (Scramble) knockdown, hairpins were cloned into Tet-pLKO-puro plasmid (21915; Addgene). Lentiviral supernatants were generated by transfecting the plasmids into 293T cells and used to infect 8988T cells that were then selected for at least 7 days with 2 μg ml^−1^ puromycin (Sigma-Aldrich, P8833). The RNAi consortium clone IDs for the hairpins used were: shGS #1 (G05) TRCN0000343992 (CACACCTGTAAACGGATAATG); shGS #2 (G06) TRCN0000344059 (ATAACCACTGCTTCCATTTAA); Control Scramble shRNA sequence (CCTAAGGTTAAGTCGCCCTCGCTCGAGCGAGGGCGACTTAACCTTAGG) was the same as that from Addgene Plasmid # 1864.

### Cell proliferation and cell death assays

Cells were seeded on day 0 in their respective maintenance culture medium in 96-well plates, at a density of 3,000 cells per well, and incubated overnight. On day 1, the cells were washed once with phosphate buffered saline (PBS) and incubated, without replenishment, in their experimental medium supplemented with propidium iodide (2 μg ml^-1^) for up to 7 days. Live or dead cells (propidium iodide-negative or -positive, respectively) were counted on the indicated days using the Celigo Image Cytometer (Nexcelom Bioscience), and either normalized to day 1 (live cell proliferation curves), or presented as percentage of propidium iodide-positive cells per total cell number (cell death).

### Clonogenic assay

600 cells were seeded per well in 6-well plates in their appropriate maintenance medium and incubated overnight. The next day, cells were washed once with PBS and incubated in their experimental medium for 7-10 days. Colonies were then either directly counted using the Celigo Image Cytometer (Nexcelom Bioscience), or first stained and then counted using the following protocol: media were removed and the cells fixed in a solution of 0.2% Crystal Violet (Sigma-Aldrich, C0775) in 80% methanol for 15-30 min. The wells were then washed in distilled water and allowed to dry overnight, before colonies were counted using the Celigo Image Cytometer.

### Sphere formation assay

50 cells were seeded per well in 96-well plates and incubated overnight in their appropriate maintenance medium supplemented with 4% matrigel (Corning, 356231). The next day, cells were washed once gently with PBS and treated with the experimental medium. Spheres were counted after 7-10 days using the Celigo Image Cytometer (Nexelcom Bioscience).

### Macropinocytosis assay

Cells were seeded onto glass coverslips (Bellco Glass, 1943-10012A) in 24-well plates and incubated overnight in their maintenance culture medium. The next day, the medium was replaced with fresh L-L medium containing 10% dialyzed FBS, and the cells were incubated for 24 h. Macropinosomes were assayed as previously described (41) and outlined below. 70-kDa TMR-dextran (Thermo Fisher Scientific, D1818) was added to the cells at a final concentration of 1 mg ml^-1^ and the cells incubated for 30 min at 37°C, in the absence or presence of 25 μM EIPA (Sigma-Aldrich; A3085). Cells were then rinsed 3 times on ice with ice-cold PBS and immediately fixed in 4% paraformaldehyde (EMS, 15710-S) for 30 min. After staining the cells with 1 μg ml^-1^ DAPI (Thermo Scientific, D3571) in PBS for 15min, the coverslips were mounted onto glass slides (VWR, 48311-703) using Fluoroshield mounting medium (Sigma, F6182). Image acquisition was performed using a Nikon Eclipse 90i Advanced Automated Research Microscope equipped with a standard optical filter set including DAPI and Rhodamine/Texas Red. Images were captured with a 40x objective and the NIS-Elements Advanced Research Microscope Imaging Software (Nikon). Exposure time for DAPI acquisition was 100 ms and that for TMR acquisition was 400 ms. Representative images were captured in at least 3 different fields per well in duplicate wells, per condition and analyzed using ImageJ software (NIH). At least 200 cells were counted per condition. Macropinosomes were visualized after setting the threshold of the “Auto Contrast” feature in Image J to 134. The number and area of labeled macropinosomes were then determined using the “Analyze Particles” feature, and macropinocytic uptake was computed by the total particle number and area per cell.

### U-^14^C-asparate incorporation into DNA

Cells were seeded overnight (57,000 per well) in 6-well-plates with triplicate wells per condition. The cells were then washed twice with PBS and incubated with either H-H or L-L medium supplemented with 1μCi per well of L-[U-^14^C]-Aspartic acid (PerkinElmer, NEC268E050UC) for 40 h. The cells were washed twice with PBS before DNA was extracted using PureLink® Genomic DNA Mini Kit (Thermo Fisher-K182001) and quantified using a spectrophotometer. Equal volumes of DNA were added to scintillation vials and radioactivity was measured by liquid scintillation counting and normalized to total DNA concentration.

### Immunoblotting

For western blotting, cells were rinsed once in ice-cold PBS and collected in lysis buffer containing 50 mM HEPES KOH, pH 7.4, 40mM NaCl, 2mM EDTA, 1.5mM orthovanadate, 50mM NaF, 10mM pyrophosphate, 10mM glycerophosphate, EDTA-free protease inhibitors (Thermo Fisher Scientific, 88266) and 1% Triton X-100. Proteins from total lysates were resolved by 8–12% SDS–PAGE, transferred to polyvinylidene difluoride (PVDF) and the blot was exposed to film.

### Immunoprecipitation and ubiquitination assay

To assess GS protein ubiquitination, cells were treated with either control vehicle DMSO or MG132 (10 μM) for 8 h. Cells were then lysed under denaturing conditions using the lysis buffer (40 mM HEPES KOH, pH 7.4, 1% SDS) containing 5 mM N-ethylmaleimide (NEM, Sigma-Aldrich, E1275) to prevent deubiquitination and boiled for 5 min. The lysates were diluted 10-fold in IP buffer (40 mM HEPES KOH pH 7.4, 50 mM NaF, 10 mM Sodium pyrophosphate, 10 mM Sodium beta-glycerophosphate, 1.5 mM Sodium Orthovanadate, 2.5 mM MgCl2, EDTA-free protease inhibitors, and 0.5% Triton X-100). IP was performed by incubating the samples with 2 μg of normal rabbit IgG control (Santa Cruz, sc-2027) or 2μg GS antibody (Abcam, ab73593) overnight with rotation at 4°C. Protein G agarose bead slurry (Invitrogen, 15920-010) was added to the pre-cleared lysates (60 μl, 50:50) and incubated with rotation for 1.5-2 hr at 4°C. The beads were then washed once with IP buffer, followed by 3 additional washes with IP buffer supplemented with 500 mM NaCl. Loading buffer was added to the immunoprecipitated proteins, which were then boiled, resolved by SDS-PAGE and transferred to a PVDF membrane. Ubiquitinated GS was identified by HRP-conjugated anti-Ubiquitin antibody (1:5,000; Enzo Life Sciences, FK2, BML-PW0150-0100).

### Quantitative real-time PCR

Total mRNA was isolated with RNA STAT-60 (Tel-Test, Cs-111) according to the manufacturer’s instructions, treated with DNase I (RNase-free, Roche Molecular Biochemicals, 04716728001) and reverse-transcribed into cDNA with random hexamers using the SuperScript II First-Strand Synthesis System (Invitrogen, 18064071). Primers listed below were validated and PCR reactions performed as previously described (42). Quantitative PCR reactions were performed in triplicates using an Applied Biosystems ViiA 7 Real-Time PCR system. PCR reactions contained cDNA resulting from reverse transcription of 25ng total RNA, 150nM of each primer and 5 μl 2X-Jump Start SYBR Green PCR Mix (Invitrogen) in10 μl total volume. Relative mRNA levels were calculated using the comparative CT method and normalized to cyclophilin. The QPCR primers used were:

*CyclophilinA* (CYPA): 5’-GGAGATGGCACAGGAGGAA-3’ and 5’-GCCCGTAGTGCTTCAGCTT-3’

*GS*: 5’-AAGAGTTGCCTGAGTGGAATTTC-3’ and 5’-AGCTTGTTAGGGTCCTTACGG-3’.

### RNA-Seq sample preparation

Cells were seeded overnight at 700,000 per 10 cm plate. The cells were then washed once with PBS and incubated in L-L medium for 24 h. RNA was isolated and purified using miRNeasy Mini Kit (Qiagen-217004). Total RNA (> 150 ng) was sent to the Broad Institute Genomics Platform for library preparation. RNA quality and insert-size was assessed by Caliper LabchipGXII producing a RQS value (equivalent to RIN). RNA quantity was determined by PicoGreen. mRNA libraries were prepared using the Illumina TruSeq Stranded mRNA kit modified for improved performance and multiplexing. Libraries were pooled and sequenced with the Illumina HiSeq 4000 using 101-cycle, pair-end settings to obtain 50 million read sequencing depth per sample.

### RNA read processing and Quantification

All samples were processed using an RNA-Seq pipeline implemented in the bcbio-nextgen project (https://bcbio-nextgen.readthedocs.io/en/latest/). Raw reads were examined for quality issues using FastQC (http://www.bioinformatics.babraham.ac.uk/projects/fastqc/) to ensure library generation and sequencing data were suitable for further analysis. Reads were aligned to the hg19 build of the human genome using STAR(43). Quality of alignments was assessed by checking for evenness of coverage, rRNA content, genomic context of alignments, complexity and other quality checks. Expression quantification was performed with Salmon(44) to identify transcript-level abundance estimates and then collapsed down to the gene-level using the R Bioconductor package tximport (45). Differential expression was performed at the gene level using the R Bioconductor package DESeq2(46). Differentially expressed genes were identified using the Wald test and significant genes were obtained using an FDR threshold of 0.05. GSEA(47) was carried out on the full expression data matrix using the gene sets collection from Kyoto Encyclopedia of Genes and Genomes (KEGG) to assess the enrichment of the “adaptation to nutrient deprivation” signature. A pairwise GSEA (v4.1.0) was performed by creating ranked lists of genes using the difference of class means to calculate fold change for log scale data between the Adapted and Non-Adapted PDA clones (Fig. S4) and *P* values were obtained from permuting the gene set (1000 permutations).

### ATAC-Seq

Cells were seeded overnight at 700,000 per 10 cm plate, washed once with PBS, and incubated in L-L medium for 24 h. The assay (48) was performed on 10^6^ cells per condition. Cells were washed twice in ice-cold PBS and resuspended in 50 μl ice-cold ATAC lysis buffer (10 mM Tris.Cl, pH 7.4, 10 mM NaCl, 3 mM MgCl_2_, 0.1% (v/v) Igepal CA-630), and centrifuged at 500 g at 4° C to isolate nuclear pellets that were treated in 50 μL reactions with Nextera Tn5 Transposase (Illumina, FC-121-1030) for 30 min at 37° C. Column-purified DNA (QIAGEN) was stored at 20° C or amplified immediately in 50 μl reactions with high-fidelity 2X PCR Master Mix (New England Biolabs) using a common forward primer and different reverse primers with unique barcodes for each sample. From the reaction mix, 45 μl was kept on ice after 5 cycles of PCR, while 5 μl was amplified by qPCR for 20 additional cycles; the remaining 45 μl was then amplified for the 5-7 cycles required to achieve 1/3 of the maximum qPCR fluorescence intensity. Amplified DNA was purified over columns and primer dimers (<100 bp) were removed using AMPure beads (Beckman Coulter). Size distribution of the amplified DNA was analyzed using High-sensitivity Qubit dsDNA Assay Kit (ThermoFisher). High throughout sequencing was performed by Novogene Corporation, Inc. (Sacramento, CA). Computational analysis was performed as described previously (49).

### Whole Exome Sequencing (WES)

Cells were seeded overnight at 700,000 per 10 cm plate, then washed once with PBS and incubated in L-L medium for 24 h. DNA was isolated and purified using Genomic DNA Mini Kit (Invitrogen, K-1820-01). Library preparation (Agilent SureSelect Human All Exon V6 kit) and high throughout sequencing (Illumina NovaSeq 6000, pair-end, 100 X coverage), were performed by Novogene Corporation, Inc. (Sacramento, CA). Variants were called in 9 WES samples: SUIT-2 non-adapted (C1-C3), adapted (C4-C6) or reverse-adapted (C4-C5) clones in a single batch using HaplotypeCaller in gatk4.1.4.1 (50) as implemented in bcbio-nextgen variant2 pipeline (https://doi.org/10.5281/zenodo.3564938). Main steps of the pipeline included: read alignment with bwa mem (51), bam file processing with samtools (52) and sambamba (53), quality control with multiqc (54), coverage analysis with mosdepth (55), variant annotation with snpEff (56) and vcfanno (57). Gnomad database was used as annotation resource for variant population frequencies (58). High and Moderate impact variants were defined according to snpEff documentation. High impact variants include stop-gain, frameshift and splice site variants. Moderate impact variants include mostly missense variants. For mTOR signaling pathway: variant calling resulted in 213,965 variants with read depth > 50 across 9 samples. Variants were filtered for High and Moderate impact, yielding 17,226 variants, 10 of which are located in the 23 genes from the mTOR pathway list in Figure S4*G*. All variants in mTOR pathway genes were shared between all 9 samples, none of the variants segregated according to adaptation phenotype (Dataset S1).

For all genes: Variants of High or Moderate impact were selected, rare in human population (gnomad AF < 1%), and absent at least in one sample. Variants present in all samples were removed. This resulted in 581 variants, 117 of which were of High impact (Dataset S2, sheets 1 and 2). High impact samples were clustered based on the presence or absence of High variants (Fig. S10*A*). Clustering was performed using R, r-studio, using ggplot2 and tidyverse packages, ggdendro, cowplot and hclust.

### Metabolite tracing, extraction and quantification

For stable isotope tracing experiments, cells were incubated for 24 h in L-L or H-H medium supplemented with either 0.8mM ^15^N-ammonium chloride (99% enrichment, NLM-467-1) or 0.4mM ^15^N-leucine (98% enrichment, NLM-142-0.5) purchased from Cambridge Isotope Laboratories. Cells were then rinsed in ice-cold PBS and metabolites extracted with ice-cold 80% methanol, vortexed for 10min at 4°C, and centrifuged at top speed (10min, 10,000 × g, 4°C). Supernatants were transferred to fresh Eppendorf tubes and dried using a SpeedVac. Dried extracts were suspended in 100 μl water and centrifuged at top speed for 10 min, and the supernatant was analyzed using LC-MS. For Figures 2*A*, 3*C*, 3*E*, and S5, data analysis and natural abundance correction of the stable isotope labeling data were performed as previously described (59). For Figures 3*F* and S9, data were acquired using a hydrophilic interaction liquid chromatography (HILIC)-positive ion mode mass spectrometry (MS) method operated on Nexera X2 UHPLC (Shimadzu Scientific Instruments, Marlborough, MA) coupled to a Q Exactive orbitrap mass spectrometer (Thermo Fisher Scientific, Waltham, MA) as described previously (60), with full scan high resolution MS data collected over 70–800 m/z. For global steady-state metabolite profiling, data were analyzed using MetaboAnalyst 4.0 (61), median-normalized, log-transformed, mean-centered and divided by the S.D. of each variable. For heatmaps, the intensity of the red or blue color varies depending on the number of groups included in a specific analysis (e.g. Fig. 2*A* and Fig. 3*E*, compared to Fig. S5*A*). This is because the intensity reflects the relative levels of each metabolite among all groups, rather than its absolute levels. Nevertheless, increased or decreased metabolite levels between any 2 groups remain evident.

### Animal work

All mouse studies and procedures were approved by the Animal Care and Use Committee at Boston Children’s Hospital. For orthotopic xenografts, 750 × 10^3^ cells suspended in 25 μl 33% Matrigel (Corning, 356231) in Hank’s Balanced Salt Solution (HBSS) were injected into the pancreata of male B6.129S7-Rag^1tm1Mom^/J mice termed Rag1^−/−^ mice (Jackson Laboratory #002216). *In vivo* ultrasound imaging (Vevo 2100, MS550D Scanhead) was performed on mice to detect and quantify tumor growth at the Small Animal Imaging Lab (SAIL) at Boston Children’s Hospital. Following experimental endpoint euthanasia, tumors were collected, their dimensions (**a, b** and **c**) were measured with a caliper and tumor volume was estimated according to the ellipsoid formula(62): 4/3 × π × (a/2 × b/2 × c/2). Tumor tissue was then either immediately frozen in liquid nitrogen, or fixed in formalin for later processing.

### Immunohistochemistry and analysis of human samples

Formalin-fixed paraffin-embedded murine PDA tumors were sectioned and stained with Hematoxylin and Eosin (H&E), Ki-67 (BD Biosciences, #550609) or cleaved caspase 3 (CST, #9664). Human PDA tissue microarrays (TMAs) with 2–3 cores per case, were sectioned and immunostained with GS antibody (abcam, ab-73593). Immunostaining was performed according to the manufacturer’s protocol. A pathologist scored, in a blinded fashion, the intensity of staining on a scale of 0-3 that represented none (0), weak (1), moderate (2), or strong (3) based on the PDA cell staining. Scores for each tumor core were estimated using the following formula: H-score = 0 × (% area of tumor with no staining) + 1 × (% tumor area with weak staining) + 2 × (% tumor area with moderate staining) + 3 × (% tumor area with strong staining), where 300 would be the highest score. TMAs were derived from 127 patients with surgically resected PDA from Massachusetts General Hospital. Informed patient consent was waived by the Institutional Review Board or IRB (Protocol Number 2009P001838), as the TMAs were generated from discarded material following clinical diagnosis. For analyses of human data, the following variables were from electronic medical records: age, sex, pN and pT stage based on the American Joint Committee on Cancer 8th edition, tumor size, location and grade of differentiation, and presence of lymphovascular and perineural invasion. Survival analyses were performed using overall survival (OS) as the outcome of interest, defined as the time between surgical resection and death from any cause or date of last clinical contact. Survival outcomes were compared based on tertiles of GS staining intensity as assessed by H-scores using the Kaplan-Meier method. Tertile thresholds were 115 and 150 H-score units. As secondary analyses, GS intensity was evaluated for its association with baseline clinicopathological characteristics at the time of surgery using the Kruskal-Wallis test. Statistical significance was set at *P* <0.05. Data analysis was performed using SAS 9.4 (version 9.4, SAS Institute, Cary, NC).

### Statistical Analysis

Data were presented as mean ± S.D. or ± S.E.M., unless otherwise indicated. When comparing two groups, a two-tailed non-paired Student’s t-test was conducted. For three or more groups, one-way ANOVA (for one factor) or two-way ANOVA (2 factors) was conducted. ANOVA was followed by post hoc Tukey’s multiple-comparison test. *p* < 0.05 was considered statistically significant.

## Acknowledgements

We thank members of the Kalaany lab and the Division of Endocrinology for helpful comments and discussions. This work was supported by the American Cancer Society Research Scholar Grant (RSG-17-070-01-TBG) to N.Y.K. and funds from Boston Children’s Hospital. N.Y.K. is also supported by NIH/NCI Grant R01 CA211944. P-Y.T. was partly funded by Top University Strategic Alliance (TUSA) fellowship from Taiwan. Work by M.M. and S.N. was supported by Harvard Catalyst, The Harvard Clinical and Translational Science Center (NIH award #UL1 RR 025758) and computations were run on the Orchestra cluster supported by HMS Research Computing Group.

## Author Contributions

N.Y.K. conceived, designed and supervised the study. P-Y.T. and M-S.L. performed all experiments with help from lab members and collaborators. M-S.L. performed biochemical assays for GS stabilization and growth assays for stable GS knockdown with help from I.N. who also assisted in macropinocytosis quantification and western blotting. A.A. assisted P-Y.T. with tumor cell injection into mice. M.M. analyzed RNA-Seq data and M-S.L. performed pathway analyses (Fig. S4). S.N. performed WES analysis. R.A.M. supervised the ATAC-Seq experiment and analyses, contributing to the interpretation of the results. U.J. performed the ATAC-Seq assay and analyzed the ATAC-Seq and RNA-Seq data (Fig. 4*E-G* and Fig. S10*B* and *C*) with help from S.M. C.A.L. performed metabolite quantification and tracing analyses (Figs 2*A* and 3*C* and *E* and Fig. S5). D.S.H., with supervision from C.B.C., performed metabolite quantification and tracing analyses (Fig. *3F* and Fig. S9). F.R.R. assisted with mouse ultrasound imaging. M.M.-K. provided TMAs and pathological assessment and scoring for immunostained human pancreatic tissue. T.H. and K.C.H. helped retrieve patient data. P.D. performed the statistical analyses on human data with consultation from V.M.-O. who contributed to the discussion of the results. N.Y.K., P.-Y.T. and M-S.L. wrote the manuscript with feedback from all authors.

## Supplementary information

Supplementary Figures S1-13 and Datasets S1 and S2 accompany this manuscript.

## Data availability

The RNA-Seq, ATAC-Seq and DNA-Seq (WES) data have been deposited in NCBI’s Gene Expression Omnibus and are accessible through GEO Series accession number GSE144833.

## Competing interests

The authors declare no competing financial interests.

## Notes

### Competing Interest Statement

The authors have declared no competing interest.

### Summary of Updates

Whole exome sequencing (WES) analysis of all clones described in this manuscript revealed an inadvertent mistake in the identity of the adapted clones from one of the human pancreatic cancer cell lines used, PA-TU-8988T. As demonstrated in the revisions, the conclusions of the study were not affected. We found that 8988T adapted (but not non-adapted) clones were actually derived from the second cell line used, SUIT-2. To correct this, all figures and figure panels containing data from 8988T adapted and non-adapted clones were removed and replaced with newly generated data derived from newly adapted and non-adapted clones of the authenticated 8988T cell line. These include Figures 1C-F (left panels), 2A, 2C (left panel), 3B (right panel), 3E, 4E, S2A-F (left panels), S2G, S5A and B, S6A (left panel), S7B (right panel). Moreover, Figures 6A-E and S12A-F have been corrected to indicate that the adapted clones used in this experiment were from SUIT-2 cells (not 8988T), as authenticated by DNA sequencing (WES) and short tandem repeat profiling (STR). In addition, several figures, figure panels and 2 Datasets have been added, representing additional experiments and analyses that further strengthen the conclusions of our findings. These include Figure 1B, 1G and 1H, Supplementary Figures S1A-C, S4A-F, S6B, S8A-D, S9, S10A, S12G and S12H and Supplementary Datasets 1 and 2. All sequencing data have been deposited in Gene Expression Omnibus database and are accessible through GEO Series accession number GSE144833.

