## Supplementary Figures and Legends for "Adaptation of pancreatic cancer cells to nutrient deprivation is reversible and requires glutamine synthetase stabilization by mTORC1"

**Supplementary Information includes 13 Supplementary Figures S1-S13 and 2 Datasets S1 and S2.**

#### Supplementary Figure Legends

**Figure S1. Culture of PDA cells in low glucose-low glutamine medium results in selection of rare surviving cells.** (*A* and *B*) Percent cell death (*A*) and relative live cell number (*B*) in 7 human PDA cell lines (SUIT-2; 8988T; MIA PaCa-2; PANC-1; BxPC-3; AsPC-1, HPAC) grown for 65 days in L-L medium (n=3). (*C*) Table representing genetic alterations in the 7 PDA cell lines described in *A*. Data represent the mean ± S.D. in *A* and *B*. *$p < 0.05$; **$p < 0.01$; ***$p < 0.001$; ****$p < 0.0001$, two-way ANOVA followed by Tukey test. In *A*, stars indicate statistical significance on the indicated days and Day 1 for each cell line; In *B*, stars indicate statistical significance between Day 65 for each cell line that adapted and proliferated past week 2 (i.e. SUIT-2; 8988T or MIA PaCa-2) and Day 65 for all other cell lines that did not proliferate past week 2 (PANC-1; BxPC-3; AsPC-1 or HPAC). In *C*, "#" indicates non-sense mutation; "si" indicates silent mutation.

**Figure S2. Adaptation to nutrient deprivation enhances PDA cell growth potential.** (*A*) Proliferation curves of adapted SUIT-2 and 8988T PDA clones (A-C4, -C5, -C6) and control non-adapted (NA-C1, -C2, -C3) clones that were grown in H-H medium for 7 days (n=6 for SUIT-2 and n=8 for 8988T). (*B*) Percent cell death in clones described in *A* that was quantified on Day 4 (SUIT-2, n=6) or Day 5 (8988T, n=8) of growth in H-H medium. (*C*) Colony formation assay showing number of colonies formed by cells in *A* that were grown for 7 days in

H-H conditions (n=6 for SUIT-2 and n=3 for 8988T). (*D-F*) 3D growth assay showing number (*D* and *E*) or area (*F*) of spheres formed by cells in *A* that were grown for 7 days (SUIT-2) or 10 days (8988T) in L-L (*D*) or H-H (*E* and *F*) medium supplemented with 4% matrigel (n=6 and n=8 for 8988T). (*G*) Representative images of mice bearing orthotopic transplants of adapted or non-adapted SUIT-2 PDA clones. Boxed areas are 3-fold enlarged on the right, demonstrating the presence of a large pancreatic tumor derived from adapted clonal cells (A-C5) and absence of tumors on the pancreas that was previously injected with non-adapted cells (NA-C3). Data represent the mean ± S.E.M. in *A*, and mean ± S.D. in *B-F*. *$p < 0.05$; **$p < 0.01$; ***$p < 0.001$; ****$p < 0.0001$, two-way ANOVA for *A* and one-way ANOVA for *B-F*, followed by Tukey test.

**Figure S3. Autophagy and macropinocytosis are not further induced in PDA cells upon adaptation to nutrient deprivation.** (*A*) Immunoblots of the autophagy marker LC3B-II in SUIT-2 non-adapted “NA” or adapted “A” clones treated under L-L medium, with the autophagy inhibitor chloroquine (CQ, 10 μM) for 2, 6 or 24 h. Blots demonstrate similar or decreased autophagic flux in adapted compared to non-adapted cells. (*B*) Representative images for macropinocytic uptake of TMR-dextran by non-adapted or adapted SUIT-2 clonal cells in the presence or absence of the inhibitor EIPA (top) with quantification of macropinocytic area per cell (bottom). Scale bar, 25 μm. Data represent the mean ± S.D. for n=2 independent experiments with approximately 285 cells scored per experiment. *$p < 0.05$, one-way ANOVA followed by Tukey test.

**Figure S4. Adaptation to nutrient deprivation enriches for pathways involved in nucleotide and amino acid metabolism.** (*A-G*) Heatmaps of genes from enriched metabolic pathways in adapted "A" compared to non-adapted "NA" SUIT-2 clones (n = 3 per group), with corresponding GSEA plots. Represented are pathways for: *A*, Purine Metabolism; *B*, Pyrimidine Metabolism; *C*, Branched Chain Amino Acid (BCAA) Metabolism; *D*, Tyrosine Metabolism; *E*, Cysteine and Methionine Metabolism; *F*, Arginine and Proline Metabolism; *G*, mTOR Signaling. Red indicates higher expression, and blue lower expression, relative to the mean expression level for each metabolic gene across all groups. NES normalized enrichment scores, FDR false discovery rate, Nom. Nominal.

**Figure S5. Adapted but not non-adapted PDA clones treated with medium low in glucose and glutamine display a metabolite profile closer to that of clones under nutrient-replete conditions.** (*A*) Heatmap listing in descending order of statistical significance ($p < 0.05$ by $t$ test), amino acids, metabolites in the nucleotide synthesis pathway, and metabolites in the TCA cycle, glycolysis/gluconeogenesis pathways in non-adapted or adapted PDA SUIT-2 clones that were treated for 24 h with H-H or L-L medium. Data represent 3 clones per condition with n=4 replicates per clone. Red indicates higher level and blue lower level, relative to the median for each metabolite across all groups. (*B*) List of top 32 metabolic pathways in descending order of statistical significance (-log $p \geq 6.28$; FDR $\leq 2.92 \times 10^{-3}$) that are enriched in adapted, compared to non-adapted PDA clones, under L-L medium. In bold are pathways involving metabolites in *A*. FDR, false discovery rate.

**Figure S6. Phosphorylation of AKT or ERK is not increased in adapted PDA clones under nutrient deprivation.** (*A*) Immunoblots of pS473-AKT, pT389-AKT, pT202/Y204 ERK1/2, total AKT and total ERK in non-adapted "NA" or adapted "A" SUIT-2 and 8988T clones under

H-H and L-L medium. GAPDH was used as loading control. (*B*) Relative number of live cells from SUIT-2 non-adapted "NA" or adapted "A" clones treated for 48 h with increasing concentrations of rapamycin (0, 1, 10 or 20 μM; n=8 per clone per condition) in L-L medium. $*p < 0.05$; $**p < 0.01$; $****p < 0.0001$, two-way ANOVA followed by Tukey test. Stars indicate statistical significance between NA and A clones treated with the same concentration of rapamycin.

**Figure S7. Transaminase expression is induced in adapted PDA cells under nutrient deprivation.** (*A*) Relative mRNA levels of GS in adapted "A" (C4-C6) or non-adapted "NA" (C1-C3) SUIT-2 clones treated for 24 h with either H-H or L-L medium. Data represent the mean of 3 clones per condition ± S.D., n = 3 replicates per clone. $**p < 0.01$; $***p < 0.001$, two-way ANOVA followed by Tukey test. (*B*) Protein levels of transaminases BCAT1/2 and GOT1/2 in cells described in *A* treated for 24h with L-L medium. β-actin was used as loading control.

**Figure S8. Glutamine deprivation provides higher proliferative fitness to PDA cells than glucose deprivation and correlates with higher induction of glutamine synthetase** (*A*) Proliferation curves of SUIT-2 and 8988T PDA parental cell lines that were first adapted in either low glucose-high glutamine (L-H), high glucose-low glutamine (H-L) or L-L medium or non-adapted (kept in H-H medium) and then grown in either H-H (top panels) or L-L (bottom panels) medium for 7 days (n=8). (*B*) Protein levels of GS, pT389 S6K and total S6K in cells described in *A* treated for 24 h with H-H or L-L media. (*C*) Protein levels of GS, pT389 S6K and total S6K in SUIT-2 non-adapted (NA, C1-C3) or adapted (A, C4-6) clones that were treated with H-H, L-H, H-L or L-L medium for 24h. Adapted clone 5 (A-C5) treated with H-H was rerun on both gels (last lane on the right) so as to allow for normalization of loading before quantification in *D*. (*D*) Quantification by Image J software of western blot bands in *C* representing protein levels of GS and pT389 S6K that were first normalized to β-actin in each gel, and then normalized for loading between gels using A-C5 (H-H) bands in the last lane of each gel. ***$p < 0.001$; ****$p < 0.0001$, two-way ANOVA followed by Tukey test. In *A*, stars indicate significance between each adapted cell line and control non-adapted cells on Days 5 and 7. In *B* and *C*, β-actin was used as loading control.

**Figure S9. Deprivation of glutamine alone rather than glucose alone, results in metabolic alterations that more closely mimic those under deprivation of both.** Heatmap listing the top 100 polar metabolites by clustering algorithm (Metaboanalyst 4.0) in adapted PDA SUIT-2 clones that were treated for 24 h with H-H, L-H, H-L or L-L medium. Data represent 5 clones per condition with n=4 replicates per clone. Red indicates higher level and blue lower level, relative to the median for each metabolite across all groups. Stars indicate metabolites involved glucose metabolism.

**Figure S10. Reverse adaptation of PDA cells to nutrient deprivation results in partial decrease in proliferative fitness and infrequent reversal of the adaptation-induced chromatin accessibility changes.** (*A*) Whole Exome Sequencing (WES) clustering analysis (top) and High impact variant analysis (bottom) of SUIT-2 non-adapted (C1-C3) and adapted or reverse-adapted clones (C4-C6). High impact variants include stop-gain, frameshift, and splice site variants (see Methods). (*B*) Gains and losses in chromatin accessibility during adaptation (NA, non-adapted C1 and C2; A, adapted C4 and C5) of SUIT-2 clones from Fig. 4*F*, revealing significant correlation (BETA analysis) with upregulation and downregulation, respectively, of gene expression. (*C*) Representative loci showing concordant changes in ATAC-seq and RNA-seq signals at *POLE2* and *RRM2*. Dotted boxes demarcate enhancer elements linked to genes that show gain or loss of chromatin accessibility and concordant changes in gene expression in cells adapted to nutrient deprivation. The majority of enhancers that gained chromatin accessibility upon adaptation remained accessible upon reverse adaptation (e.g. *POLE2*) with some exceptions (e.g. *RRM2*). In both cases however, reversal of mRNA expression changes was partial (e.g. *POLE2* and *RRM2*).

**Figure S11. Glutamine synthetase levels are decreased post-translationally with Torin 1 treatment.** (*A*) Relative mRNA levels of GS in SUIT-2 adapted clones (C4, C5, C6) treated with L-L medium in the absence or presence of Torin 1 (200 nM) for 24 h. Data represent the mean ± S.D. (n=3 replicates per clone per condition). $^{**}p < 0.01$; $^{***}p < 0.001$; $^{****}p < 0.0001$, Student's t-test. (*B*) Immunoblots of GS, pT389-S6K and total S6K in cells from *A* that were treated under L-L medium with control DMSO or Torin 1 (200 nM) for a total of 8 h, in the absence or presence of cycloheximide (CHX, 20 $\mu g\ ml^{-1}$) for the indicated times. β-actin was used as loading control.

**Figure S12. Inhibition of glutamine synthetase suppresses the growth of PDA cells adapted to nutrient deprivation.** (*A*) Fold change in cell number of SUIT-2 adapted clones, 72 h following treatment with the GS inhibitor (MSO) compared to vehicle control (water), showing decrease in cell number upon GS inhibition under L-L but not H-H conditions (n=5 per condition). (*B*) Proliferation curves of adapted SUIT-2 clones transfected with control nontargeting siRNA or siGS, that were grown in L-L medium for 4 days (n=6). (*C*) Colony formation assay for cells described in *B* that were grown for 8 days, under L-L conditions (n=6). (*D*) Immunoblots showing GS levels in cells from *B*, 24h post-transfection with siRNA. β-actin was used as loading control. (*E*) Volumes quantified by ultrasound on day 29, of orthotopic PDA xenograft tumors derived from SUIT-2 clonal adapted cells stably expressing Doxycycline (Dox)-inducible hairpins for GS or control Scrambled hairpin. Cells (750 x $10^3$) were injected into the pancreas of 4-6 week old Rag1$^{-/-}$ mice and Dox was supplemented in the drinking water (2 mg ml$^{-1}$) 5 days later (n=5 shScr -Dox; n=7 shScr + Dox; n=8 shGS#1; n=7 shGS#2). (*F*) Representative ultrasound images on day 29 of adapted SUIT-2 orthotopic xenograft tumors described in *E*. (*G*) Representative H&E staining and immunohistochemical analyses of Ki-67 and cleaved casapase-3 in sequential sections of orthotopic PDA tumors described in *E*, that were derived from Dox-treated mice and harvested on day 33 post-tumor cell injection. (*H*) Quantification of Ki-67-positive and cleaved caspase-3-positive cells in orthotopic PDA tumors described in *G*. Data represent the mean ± S.D. in *A*, *C, E*, *H* and mean ± S.E.M. in *B*. **$p < 0.01$; ***$p < 0.001$; ****$p < 0.0001$, t-test for *C*, two-way ANOVA for *B* and *E* and one-way ANOVA for *H* followed by Tukey test.

**Figure S13. Glutamine synthetase average protein levels do not associate with human PDA tumor grade or patient survival.** (*A*) Kruskal-Wallis test for associations between GS level (H-score) and tumor size (pT), lymph node involvement (pN), or grade (tumor differentiation level) at time of resection. Low grade indicates well-differentiated tumors; Intermediate grade, moderately differentiated tumors; High grade, poorly differentiated tumors. (*B*) Representative images of human normal pancreatic parenchyma with GS staining in acinar cells (red arrowhead), duct cells (black arrowhead), islets (black arrows) and acinar ductal metaplasia (ADM, circled). Staining in human PDA shows no correlation of GS with tumor grade, as many lower-grade tumors display high GS protein levels and higher-grade tumors display low GS levels. (*C*) Kaplan Meier survival curves stratified by GS level, categorized as tertiles. Follow-up period is indicated in months. Median survival is slightly shorter for patients with low GS compared to patients with medium or high GS levels. However, the differences between the survival curves are not statistically significant, based on the Log-Rank test ($P = 0.18$). These results are consistent with a multivariate Cox proportional hazard regression model that adjusted for age, sex, and tumor grade.

**Dataset S1. Lack of High or Moderate impact mutations in mTOR signaling pathway in PDA clones upon adaptation or reverse adaptation.** This file lists 10 High and Moderate impact variants found in the 23 genes listed in Figure S4*G* in the mTOR signaling pathway that was transcriptionally enriched in PDA cells upon adaptation. All 10 variants are present in all clones, independent of adaptation. High impact variants include stop-gain, frameshift, and splice site variants. Moderate impact variants include mostly missense variants (see Methods).

**Dataset S2. Minimal differences in High and Moderate impact variants between adapted and reverse-adapted PDA clones.** This file lists all High impact-only variants (first sheet) as well as all High and Moderate impact variants (second sheet) identified in all genes in PDA clones. No High impact mutations were found, and minimal Moderate impact mutations were identified between adapted and reverse-adapted clones. High impact variants include stop-gain, frameshift, and splice site variants. Moderate impact variants include mostly missense variants (see Methods).

### Figure S1

A

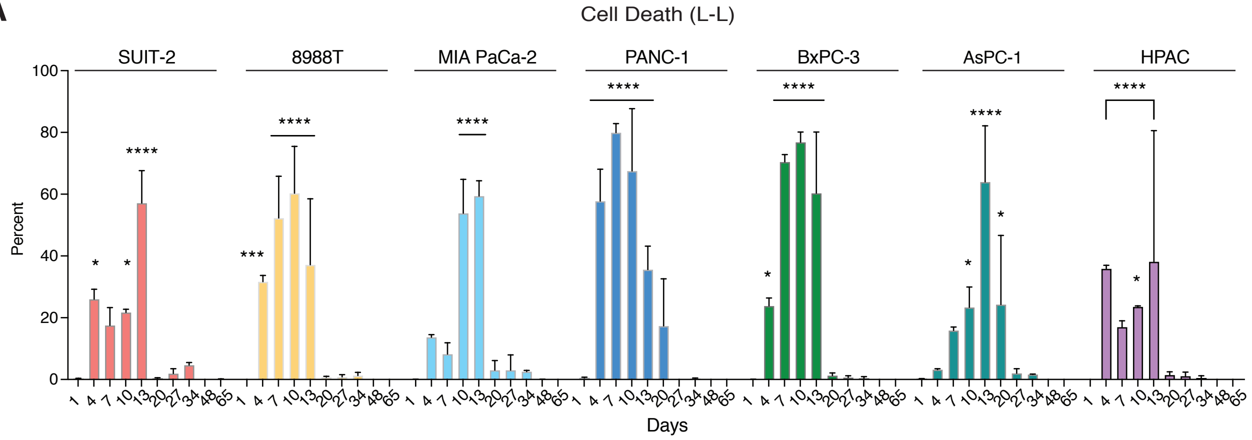


B

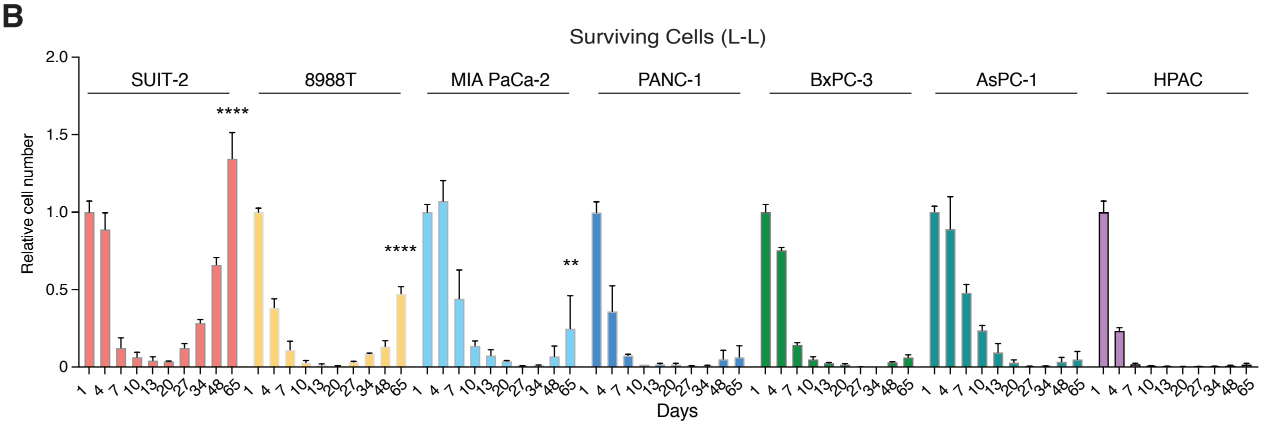


C

| Cell lines | KRAS | TP53 | CDKN2A | SMAD4 | PIK3CA | GNAS | STK11 | TGFA | BRCA1 | ALK |
|---|---|---|---|---|---|---|---|---|---|---|
| AsPC-1 | G12D | DEL | DEL | R100T | WT | A509V | WT | WT | WT | WT |
| BxPC-3 | WT | Y220C | WT | WT | WT | WT | WT | WT | WT | WT |
| HPAC | G12D | G187R | E120# | Insertion | E259D | WT | WT | WT | WT | WT |
| MIA PaCa-2 | G12C | R248W | WT | WT | WT | WT | WT | WT | WT | WT |
| PANC-1 | G12D | R273H | WT | WT | WT | WT | WT | WT | WT | WT |
| Suit-2 | G12D | R273H | E69* | WT | WT | WT | WT | WT | WT | Y406si |
| PA-TU-8988T | G12V | R282W | WT | WT | WT | WT | WT | WT | WT | WT |

#: nonsense; si: silent

#### Figure S2

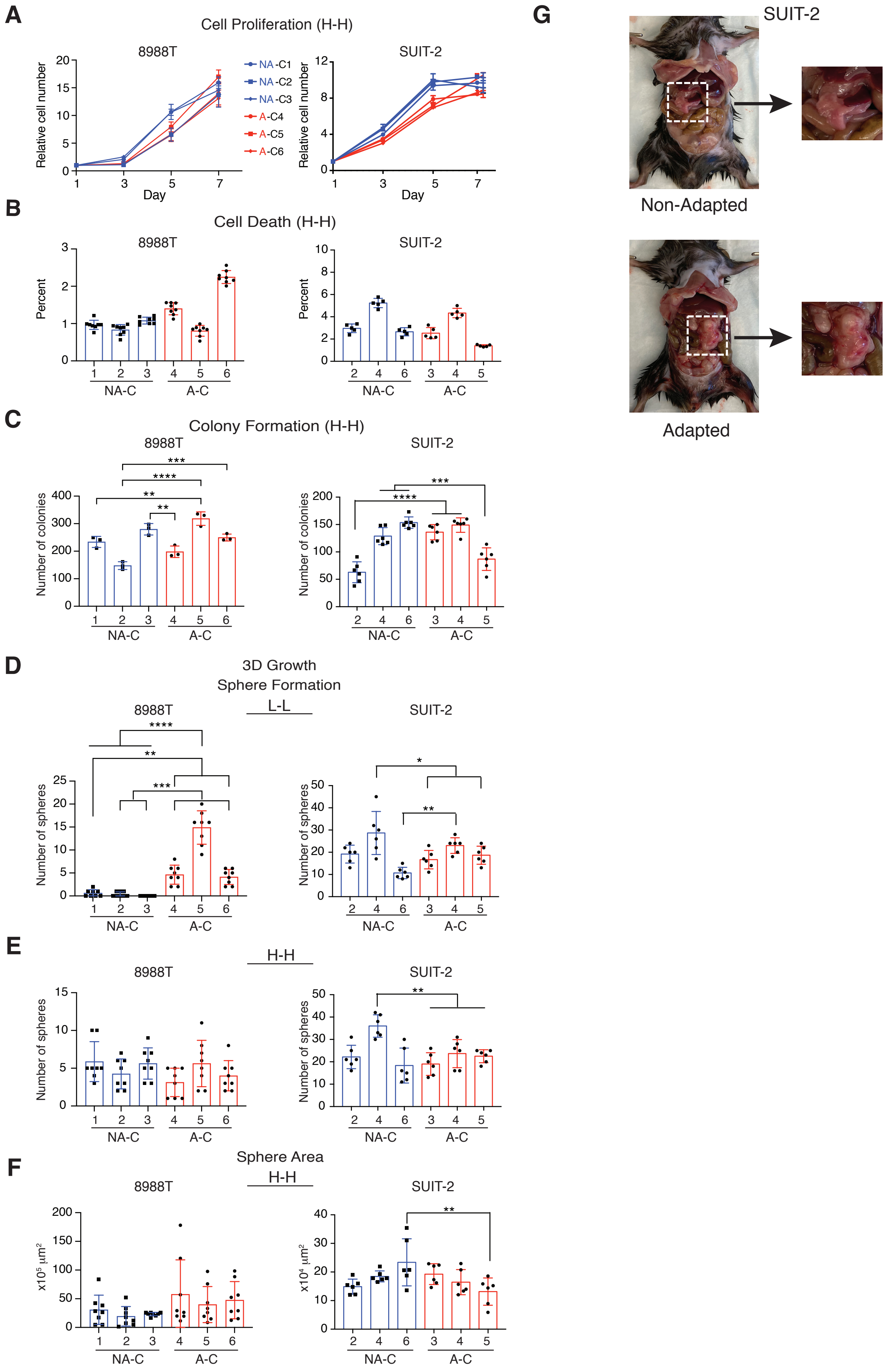

A
Cell Proliferation (H-H)
8988T
SUIT-2
Relative cell number
Day
NA-C1
NA-C2
NA-C3
A-C4
A-C5
A-C6
B
Cell Death (H-H)
Percent
NA-C
A-C
C
Colony Formation (H-H)
Number of colonies
D
3D Growth
Sphere Formation
L-L
Number of spheres
E
H-H
F
Sphere Area
x10⁵ µm²
x10⁴ µm²
G
SUIT-2
Non-Adapted
Adapted

### Figure S3

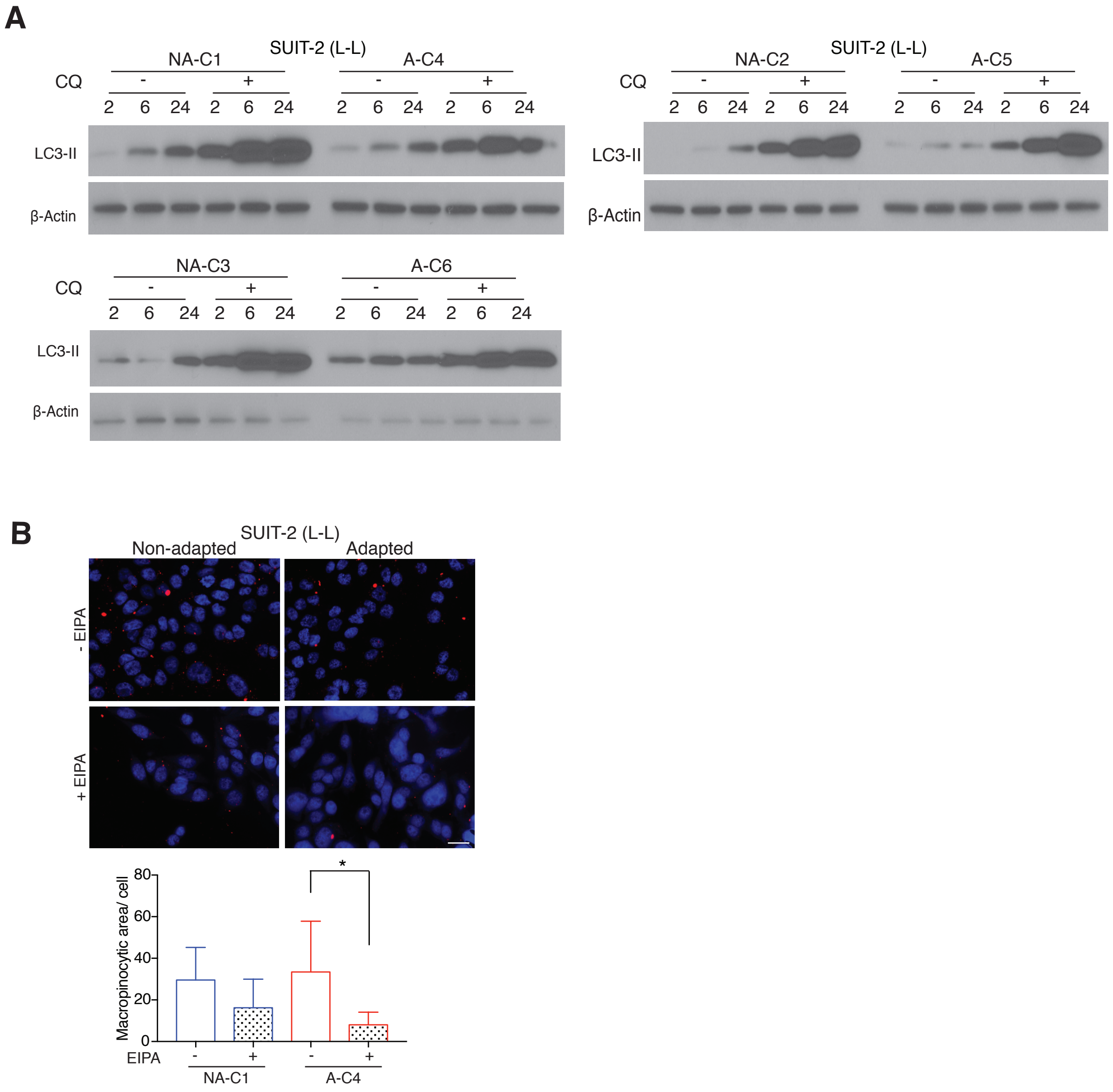
A
SUIT-2 (L-L)
NA-C1
A-C4
CQ
- + - +
2 6 24 2 6 24 2 6 24 2 6 24
LC3-II
β-Actin
SUIT-2 (L-L)
NA-C2
A-C5
CQ
LC3-II
β-Actin
NA-C3
A-C6
CQ
LC3-II
β-Actin
B
SUIT-2 (L-L)
Non-adapted
Adapted
- EIPA
+ EIPA
Macropinocytic area/ cell
80
60
40
20
0
*
EIPA
- + - +
NA-C1
A-C4

### Figure S4

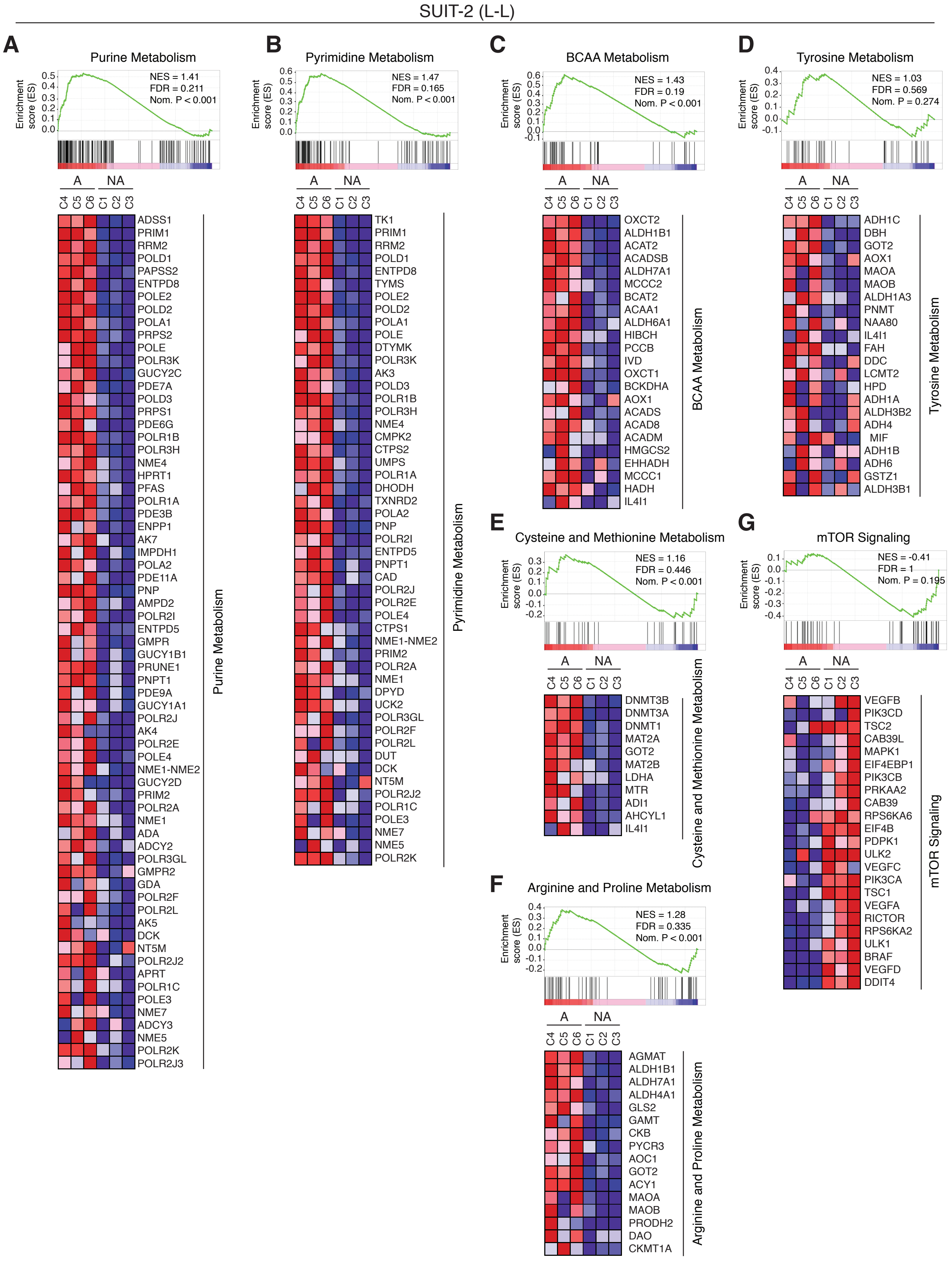
SUIT-2 (L-L)
A
Purine Metabolism
Enrichment score (ES)
NES = 1.41
FDR = 0.211
Nom. P < 0.001
A NA
C4 C5 C6 C1 C2 C3
ADSS1 PRIM1 RRM2 POLD1 PAPSS2 ENTPD8 POLE2 POLD2 POLA1 PRPS2 POLE POLR3K GUCY2C PDE7A POLD3 PRPS1 PDE6G POLR1B POLR3H NME4 HPRT1 PFAS POLR1A PDE3B ENPP1 AK7 IMPDH1 POLA2 PDE11A PNP AMPD2 POLR2I ENTPD5 GMPR GUCY1B1 PRUNE1 PNPT1 PDE9A GUCY1A1 POLR2J AK4 POLR2E POLE4 NME1-NME2 GUCY2D PRIM2 POLR2A NME1 ADA ADCY2 POLR3GL GMPR2 GDA POLR2F POLR2L AK5 DCK NT5M POLR2J2 APRT POLR1C POLE3 NME7 ADCY3 NME5 POLR2K POLR2J3
Purine Metabolism
B
Pyrimidine Metabolism
Enrichment score (ES)
NES = 1.47
FDR = 0.165
Nom. P < 0.001
A NA
C4 C5 C6 C1 C2 C3
TK1 PRIM1 RRM2 POLD1 ENTPD8 TYMS POLE2 POLD2 POLA1 POLE DTYMK POLR3K AK3 POLD3 POLR1B POLR3H NME4 CMPK2 CTPS2 UMPS POLR1A DHODH TXNRD2 POLA2 PNP POLR2I ENTPD5 PNPT1 CAD POLR2J POLR2E POLE4 CTPS1 NME1-NME2 PRIM2 POLR2A NME1 DPYD UCK2 POLR3GL POLR2F POLR2L DUT DCK NT5M POLR2J2 POLR1C POLE3 NME7 NME5 POLR2K
Pyrimidine Metabolism
C
BCAA Metabolism
Enrichment score (ES)
NES = 1.43
FDR = 0.19
Nom. P < 0.001
A NA
C4 C5 C6 C1 C2 C3
OXCT2 ALDH1B1 ACAT2 ACADSB ALDH7A1 MCCC2 BCAT2 ACAA1 ALDH6A1 HIBCH PCCB IVD OXCT1 BCKDHA AOX1 ACADS ACAD8 ACADM HMGCS2 EHHADH MCCC1 HADH IL4I1
BCAA Metabolism
D
Tyrosine Metabolism
Enrichment score (ES)
NES = 1.03
FDR = 0.569
Nom. P = 0.274
A NA
C4 C5 C6 C1 C2 C3
ADH1C DBH GOT2 AOX1 MAOA MAOB ALDH1A3 PNMT NAA80 IL4I1 FAH DDC LCMT2 HPD ADH1A ALDH3B2 ADH4 MIF ADH1B ADH6 GSTZ1 ALDH3B1
Tyrosine Metabolism
E
Cysteine and Methionine Metabolism
Enrichment score (ES)
NES = 1.16
FDR = 0.446
Nom. P < 0.001
A NA
C4 C5 C6 C1 C2 C3
DNMT3B DNMT3A DNMT1 MAT2A GOT2 MAT2B LDHA MTR ADI1 AHCYL1 IL4I1
Cysteine and Methionine Metabolism
F
Arginine and Proline Metabolism
Enrichment score (ES)
NES = 1.28
FDR = 0.335
Nom. P < 0.001
A NA
C4 C5 C6 C1 C2 C3
AGMAT ALDH1B1 ALDH7A1 ALDH4A1 GLS2 GAMT CKB PYCR3 AOC1 GOT2 ACY1 MAOA MAOB PRODH2 DAO CKMT1A
Arginine and Proline Metabolism
G
mTOR Signaling
Enrichment score (ES)
NES = -0.41
FDR = 1
Nom. P = 0.195
A NA
C4 C5 C6 C1 C2 C3
VEGFB PIK3CD TSC2 CAB39L MAPK1 EIF4EBP1 PIK3CB PRKAA2 CAB39 RPS6KA6 EIF4B PDPK1 ULK2 VEGFC PIK3CA TSC1 VEGFA RICTOR RPS6KA2 ULK1 BRAF VEGFD DDIT4
mTOR Signaling

### Figure S5

**A**

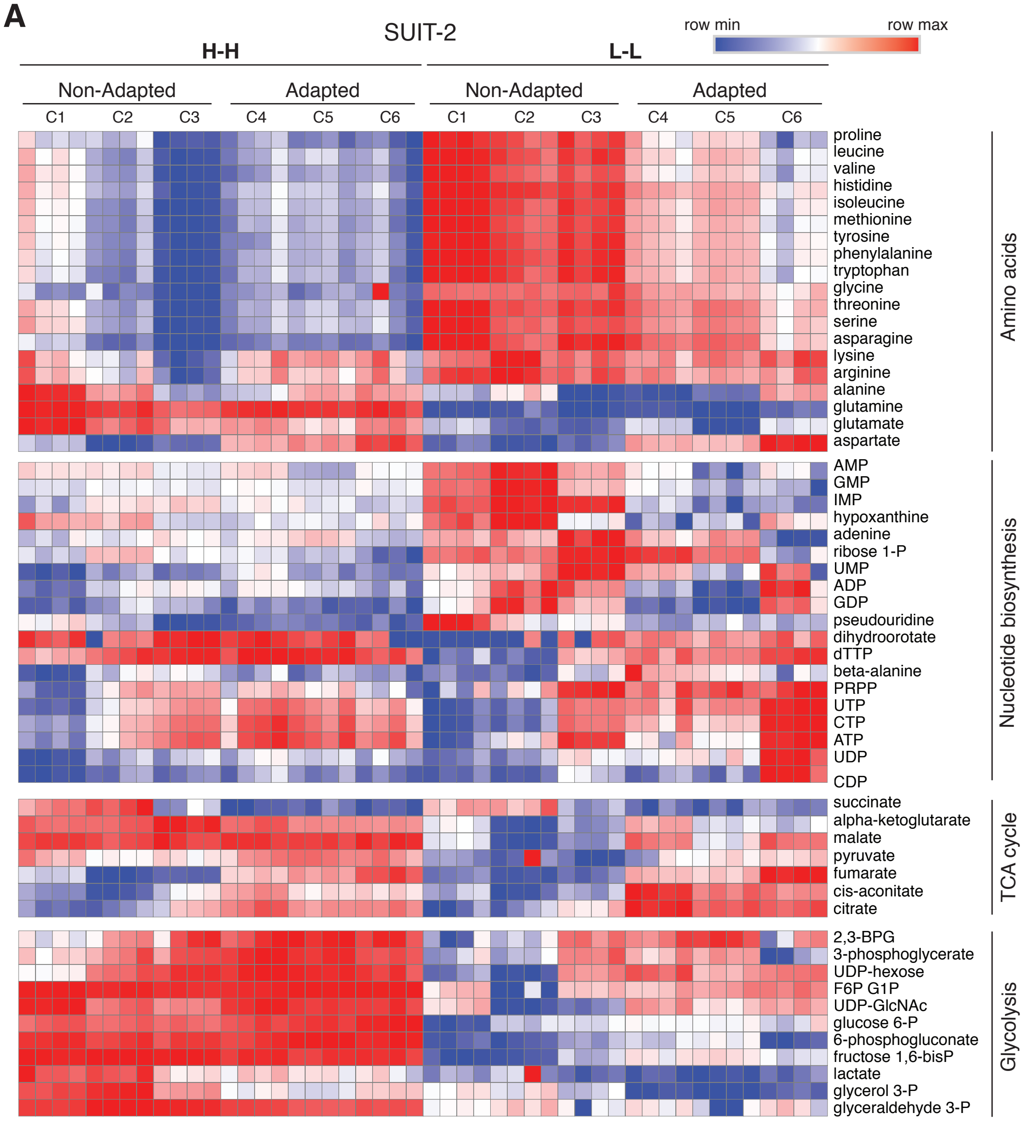


**B**

| Pathways | -log (p) | FDR |
|---|---|---|
| Glyoxylate and dicarboxylate metabolism | 2.78E+01 | 4.11E-11 |
| Alanine, aspartate and glutamate metabolism | 2.55E+01 | 1.62E-10 |
| **Citrate cycle (TCA cycle)** | **2.52E+01** | **1.62E-10** |
| Nicotinate and nicotinamide metabolism | 2.51E+01 | 1.62E-10 |
| Cysteine and methionine metabolism | 2.34E+01 | 6.59E-10 |
| Histidine metabolism | 2.31E+01 | 7.41E-10 |
| Phenylalanine, tyrosine and tryptophan biosynthesis | 2.25E+01 | 9.88E-10 |
| Phenylalanine metabolism | 2.25E+01 | 9.88E-10 |
| Glycine, serine and threonine metabolism | 2.25E+01 | 9.88E-10 |
| Ubiquinone and other terpenoid-quinone biosynthesis | 2.21E+01 | 1.24E-09 |
| Pyruvate metabolism | 2.19E+01 | 1.40E-09 |
| **Valine, leucine and isoleucine degradation** | **2.11E+01** | **2.86E-09** |
| beta-Alanine metabolism | 2.08E+01 | 3.69E-09 |
| Pantothenate and CoA biosynthesis | 2.06E+01 | 4.17E-09 |
| Tyrosine metabolism | 2.03E+01 | 5.19E-09 |
| Aminoacyl-tRNA biosynthesis | 2.01E+01 | 5.83E-09 |
| **Valine, leucine and isoleucine biosynthesis** | **1.97E+01** | **8.20E-09** |
| Arginine biosynthesis | 1.79E+01 | 4.58E-08 |
| Primary bile acid biosynthesis | 1.77E+01 | 5.49E-08 |
| Tryptophan metabolism | 1.75E+01 | 6.50E-08 |
| **Purine metabolism** | **1.73E+01** | **7.51E-08** |
| Arginine and proline metabolism | 1.66E+01 | 1.42E-07 |
| Sphingolipid metabolism | 1.63E+01 | 1.83E-07 |
| Lysine degradation | 1.33E+01 | 3.34E-06 |
| Glutathione metabolism | 1.13E+01 | 2.49E-05 |
| Glycerophospholipid metabolism | 1.11E+01 | 2.90E-05 |
| Porphyrin and chlorophyll metabolism | 1.06E+01 | 4.83E-05 |
| Propanoate metabolism | 1.04E+01 | 5.41E-05 |
| Amino sugar and nucleotide sugar metabolism | 9.01E+00 | 2.11E-04 |
| **Pyrimidine metabolism** | **8.47E+00** | **3.48E-04** |
| Butanoate metabolism | 6.83E+00 | 1.74E-03 |
| **Glycolysis / Gluconeogenesis** | **6.28E+00** | **2.92E-03** |

#### Figure S6

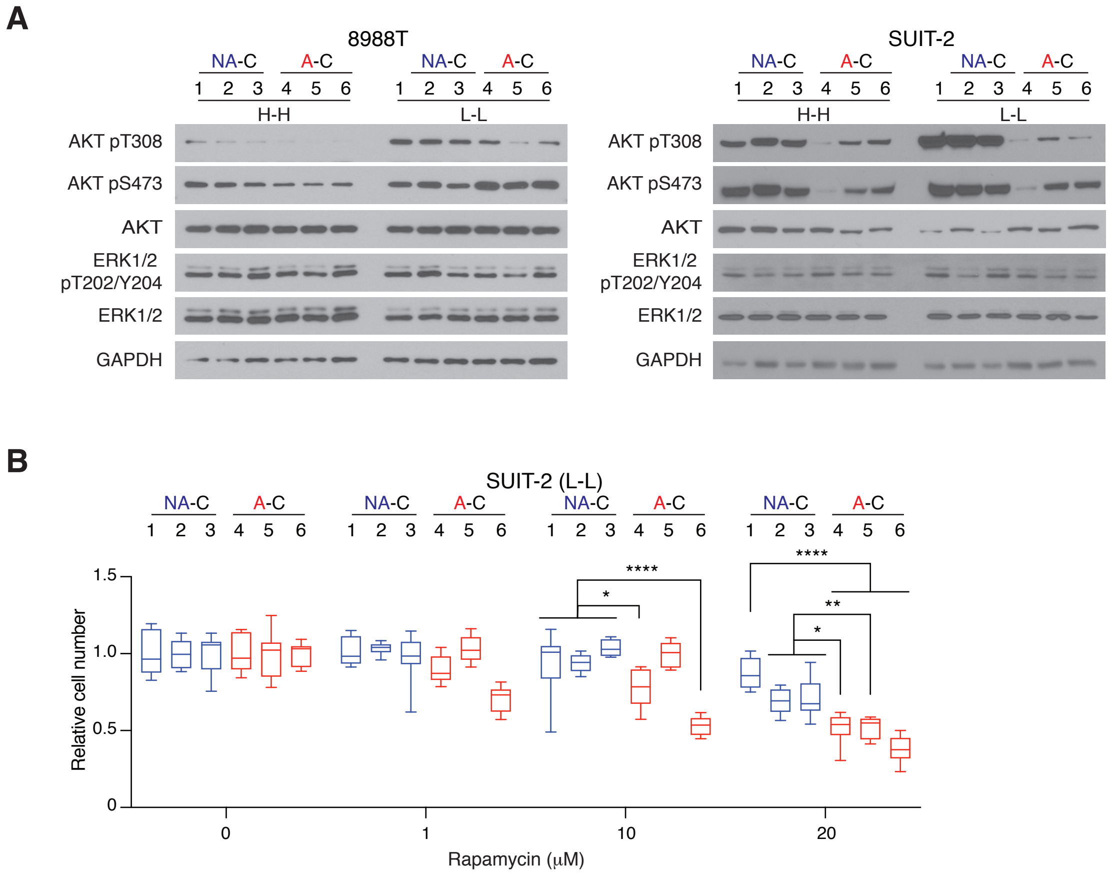
A
8988T
SUIT-2
NA-C
A-C
1 2 3 4 5 6
H-H
L-L
AKT pT308
AKT pS473
AKT
ERK1/2 pT202/Y204
ERK1/2
GAPDH
B
SUIT-2 (L-L)
Relative cell number
1.5
1.0
0.5
0
0
1
10
20
Rapamycin (μM)
****
*
**

### Figure S7

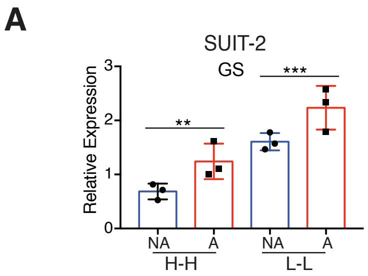
A
SUIT-2
GS
Relative Expression
3
2
1
0
**
***
NA
A
NA
A
H-H
L-L

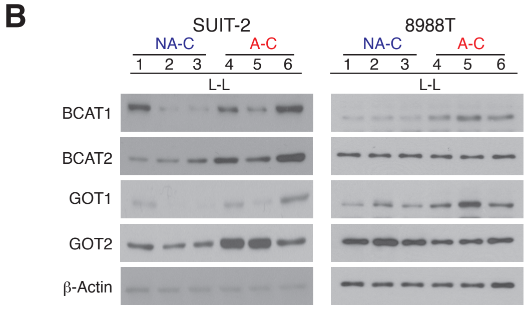
B
SUIT-2
NA-C
A-C
1 2 3 4 5 6
L-L
8988T
NA-C
A-C
1 2 3 4 5 6
L-L
BCAT1
BCAT2
GOT1
GOT2
β-Actin

### Figure S8

A

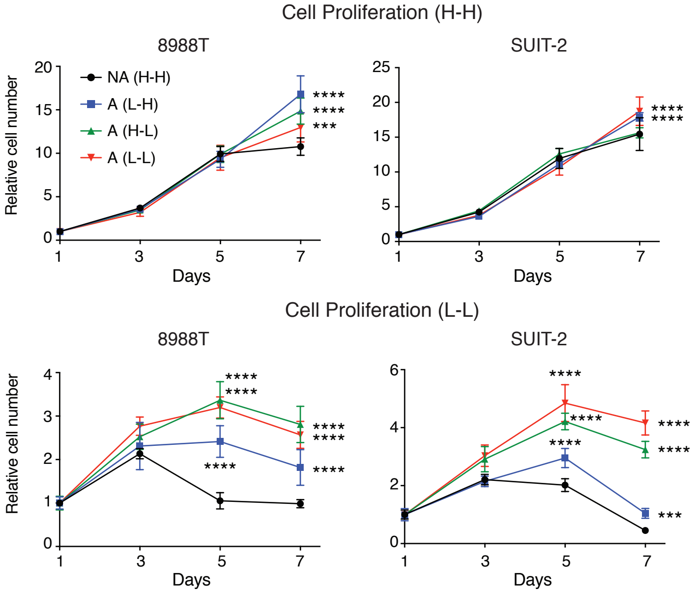


B

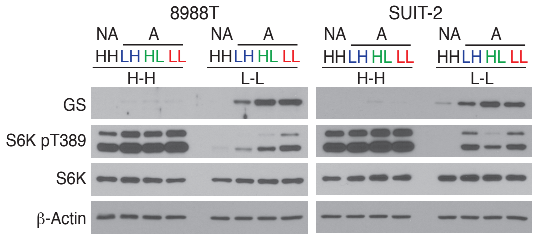


C

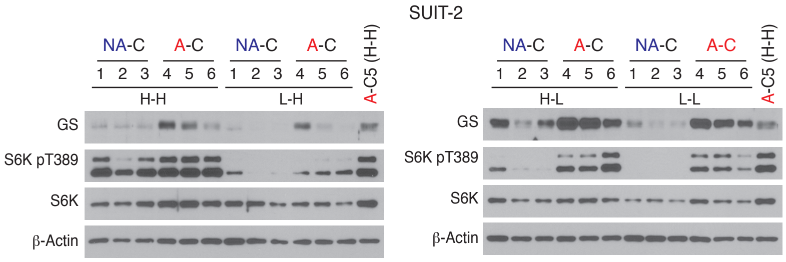


D

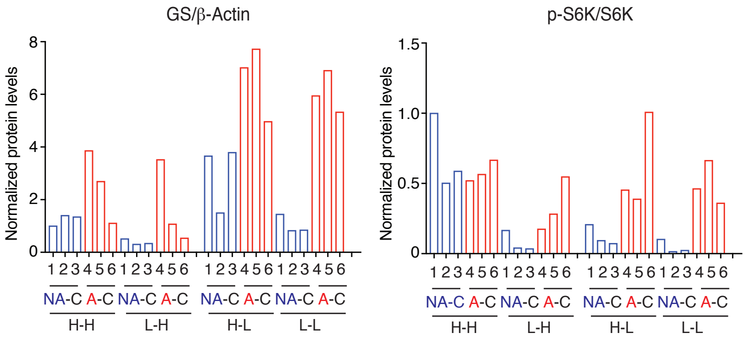

### Figure S9

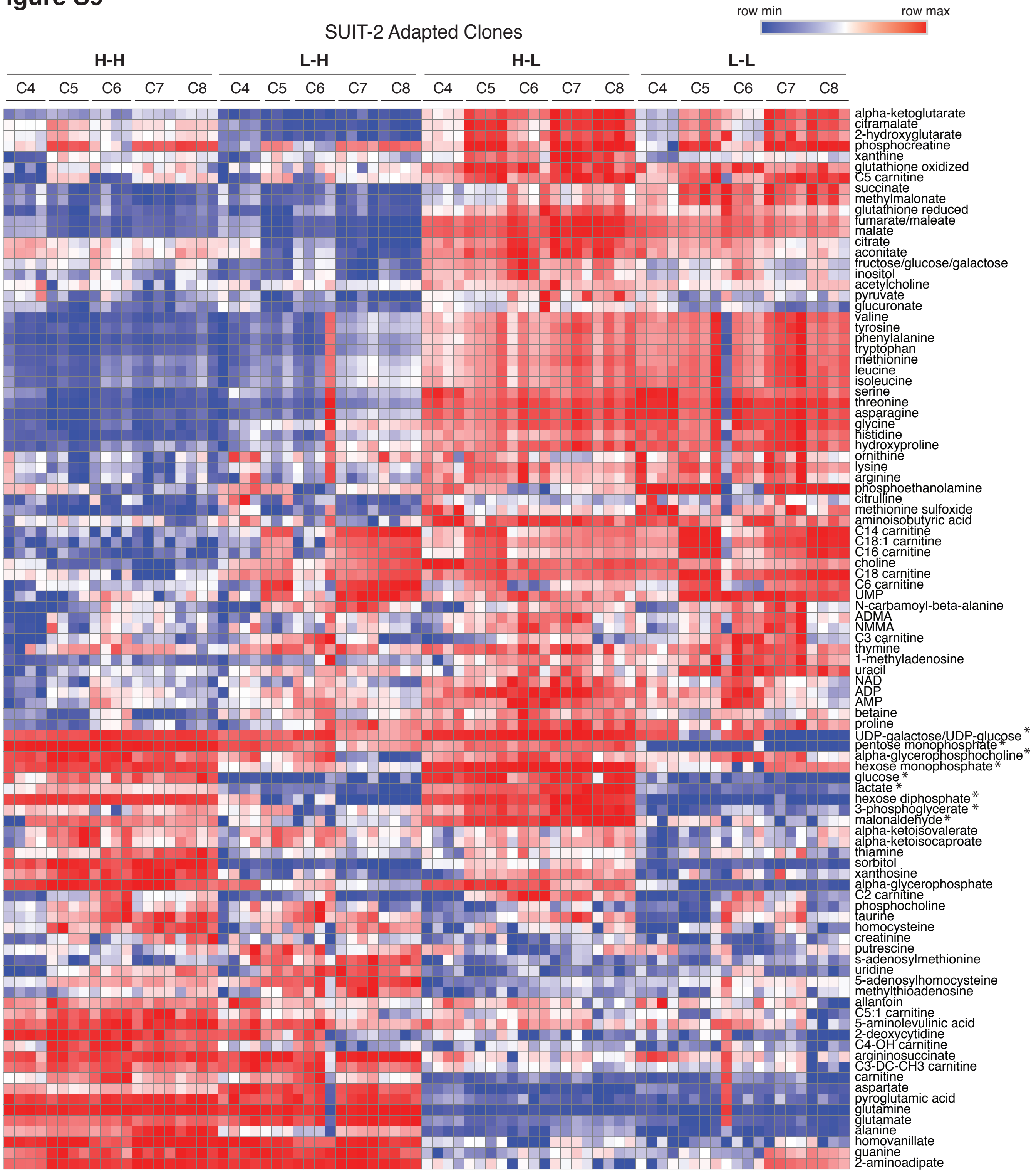

SUIT-2 Adapted Clones
row min
row max
H-H
L-H
H-L
L-L
C4
C5
C6
C7
C8
alpha-ketoglutarate
citramalate
2-hydroxyglutarate
phosphocreatine
xanthine
glutathione oxidized
C5 carnitine
succinate
methylmalonate
glutathione reduced
fumarate/maleate
malate
citrate
aconitate
fructose/glucose/galactose
inositol
acetylcholine
pyruvate
glucuronate
valine
tyrosine
phenylalanine
tryptophan
methionine
leucine
isoleucine
serine
threonine
asparagine
glycine
histidine
hydroxyproline
ornithine
lysine
arginine
phosphoethanolamine
citrulline
methionine sulfoxide
aminoisobutyric acid
C14 carnitine
C18:1 carnitine
C16 carnitine
choline
C18 carnitine
C6 carnitine
UMP
N-carbamoyl-beta-alanine
ADMA
NMMA
C3 carnitine
thymine
1-methyladenosine
uracil
NAD
ADP
AMP
betaine
proline
UDP-galactose/UDP-glucose *
pentose monophosphate *
alpha-glycerophosphocholine *
hexose monophosphate *
glucose *
lactate *
hexose diphosphate *
3-phosphoglycerate *
malonaldehyde *
alpha-ketoisovalerate
alpha-ketoisocaproate
thiamine
sorbitol
xanthosine
alpha-glycerophosphate
C2 carnitine
phosphocholine
taurine
homocysteine
creatinine
putrescine
s-adenosylmethionine
uridine
5-adenosylhomocysteine
methylthioadenosine
allantoin
C5:1 carnitine
5-aminolevulinic acid
2-deoxycytidine
C4-OH carnitine
argininosuccinate
C3-DC-CH3 carnitine
carnitine
aspartate
pyroglutamic acid
glutamine
glutamate
alanine
homovanillate
guanine
2-aminoadipate

#### Figure S10

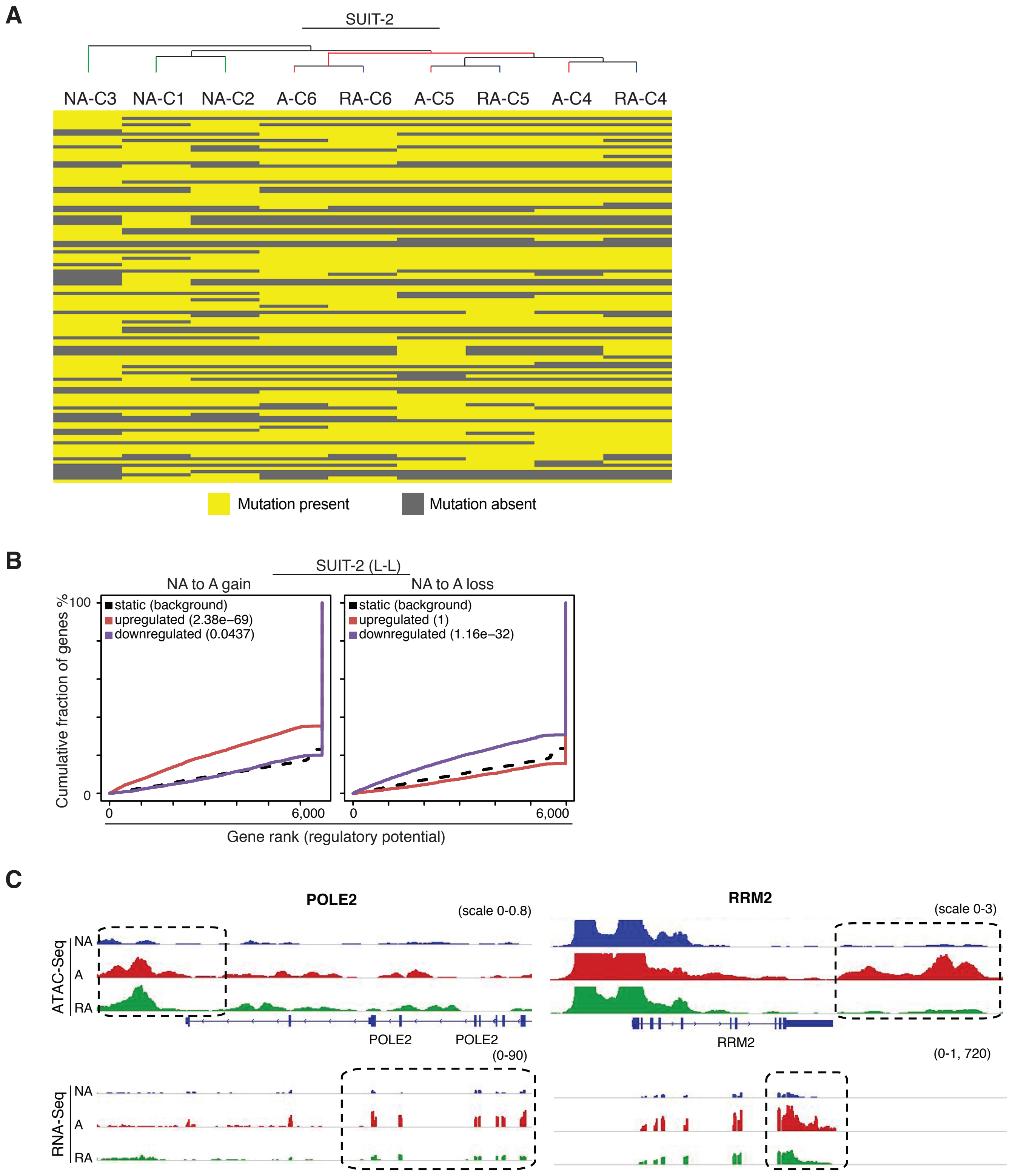
A
SUIT-2
NA-C3
NA-C1
NA-C2
A-C6
RA-C6
A-C5
RA-C5
A-C4
RA-C4
Mutation present
Mutation absent
B
SUIT-2 (L-L)
NA to A gain
NA to A loss
static (background)
upregulated (2.38e−69)
downregulated (0.0437)
static (background)
upregulated (1)
downregulated (1.16e−32)
Cumulative fraction of genes %
100
0
6,000
Gene rank (regulatory potential)
C
POLE2
(scale 0-0.8)
RRM2
(scale 0-3)
ATAC-Seq
NA
A
RA
POLE2
POLE2
RRM2
(0-90)
(0-1, 720)
RNA-Seq

### Figure S11

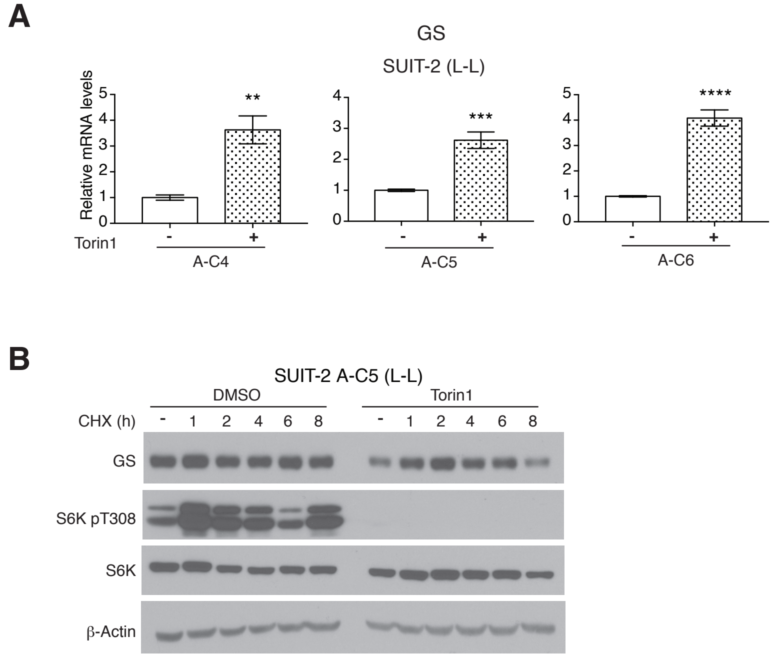

A
GS
SUIT-2 (L-L)
Relative mRNA levels
Torin1
- +
A-C4
A-C5
A-C6
**
***
****
B
SUIT-2 A-C5 (L-L)
DMSO
Torin1
CHX (h)
- 1 2 4 6 8
GS
S6K pT308
S6K
β-Actin

### Figure S12

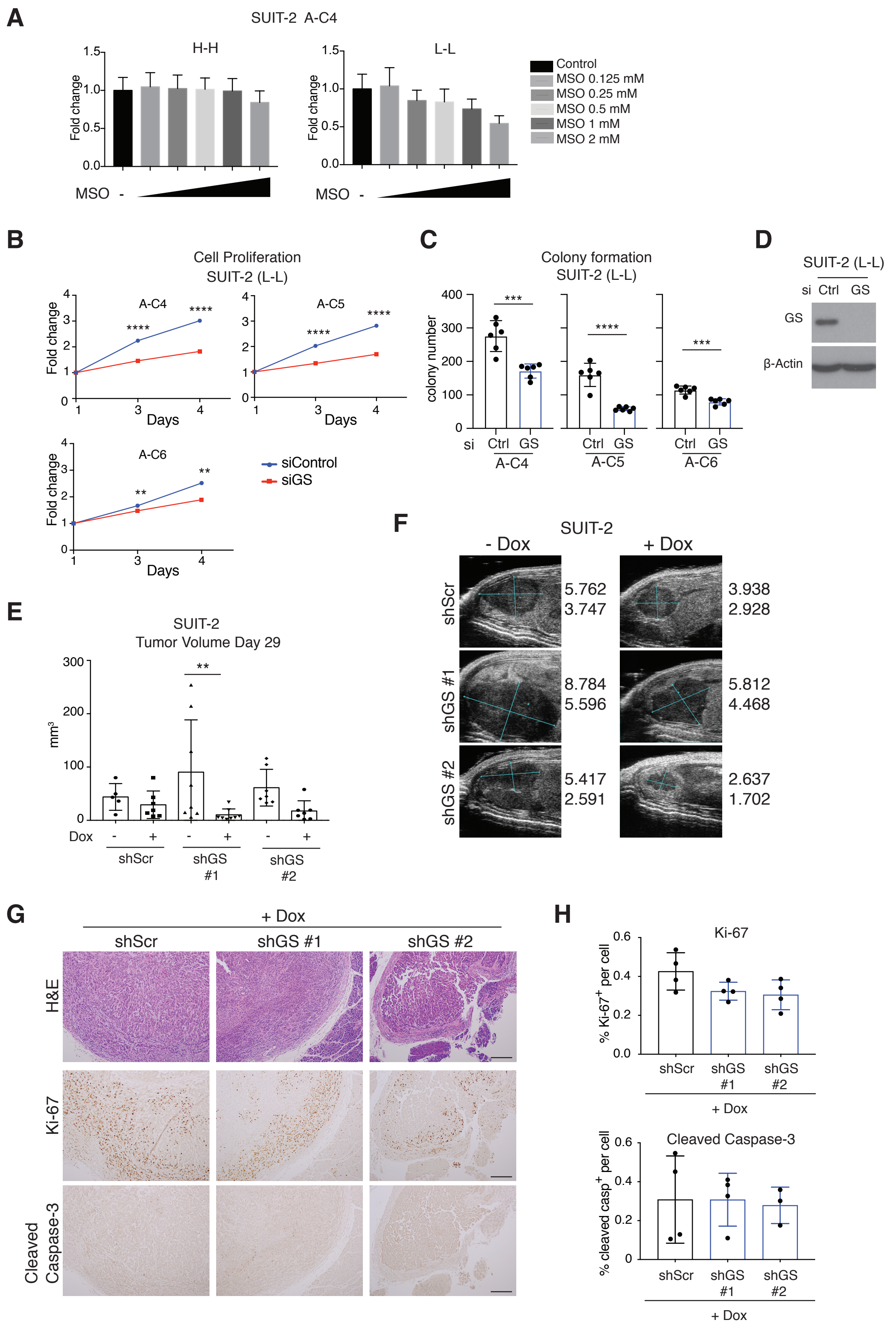
A
SUIT-2 A-C4
H-H
L-L
Fold change
MSO
Control
MSO 0.125 mM
MSO 0.25 mM
MSO 0.5 mM
MSO 1 mM
MSO 2 mM
B
Cell Proliferation
SUIT-2 (L-L)
A-C4
A-C5
A-C6
Days
siControl
siGS
C
Colony formation
SUIT-2 (L-L)
colony number
si
Ctrl GS
A-C4
A-C5
A-C6
D
SUIT-2 (L-L)
si Ctrl GS
GS
β-Actin
E
SUIT-2
Tumor Volume Day 29
mm³
Dox
shScr
shGS #1
shGS #2
F
SUIT-2
- Dox
+ Dox
shScr
shGS #1
shGS #2
5.762
3.747
3.938
2.928
8.784
5.596
5.812
4.468
5.417
2.591
2.637
1.702
G
+ Dox
shScr
shGS #1
shGS #2
H&E
Ki-67
Cleaved Caspase-3
H
Ki-67
% Ki-67+ per cell
Cleaved Caspase-3
% cleaved casp+ per cell
+ Dox

### Figure S13

**A**

TMA by pT, pN and grade

| Variable | Level | N | Median H-Score | P-value |
|---|---|---|---|---|
| pT | 1 | 14 | 150 | 0.61 |
| | 2 | 89 | 133 | |
| | 3 | 24 | 150 | |
| pN | 0 | 32 | 150 | 0.34 |
| | 1 | 62 | 125 | |
| | 2 | 33 | 150 | |
| Grade | Low | 6 | 195 | 0.32 |
| | Intermediate | 67 | 138 | |
| | High | 52 | 142 | |

**B**

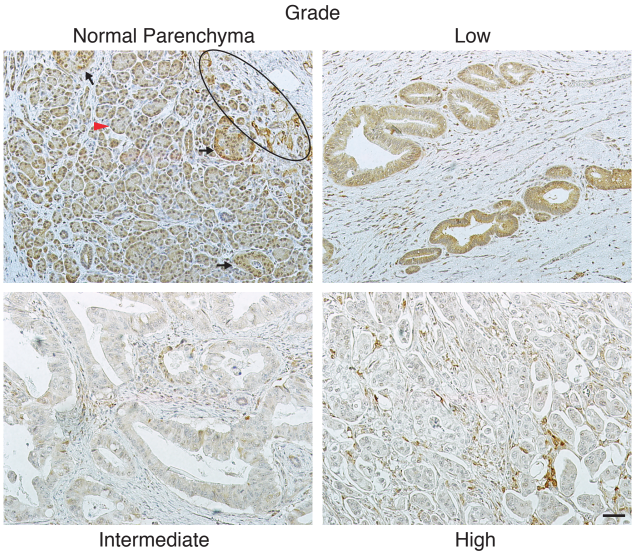


**C**

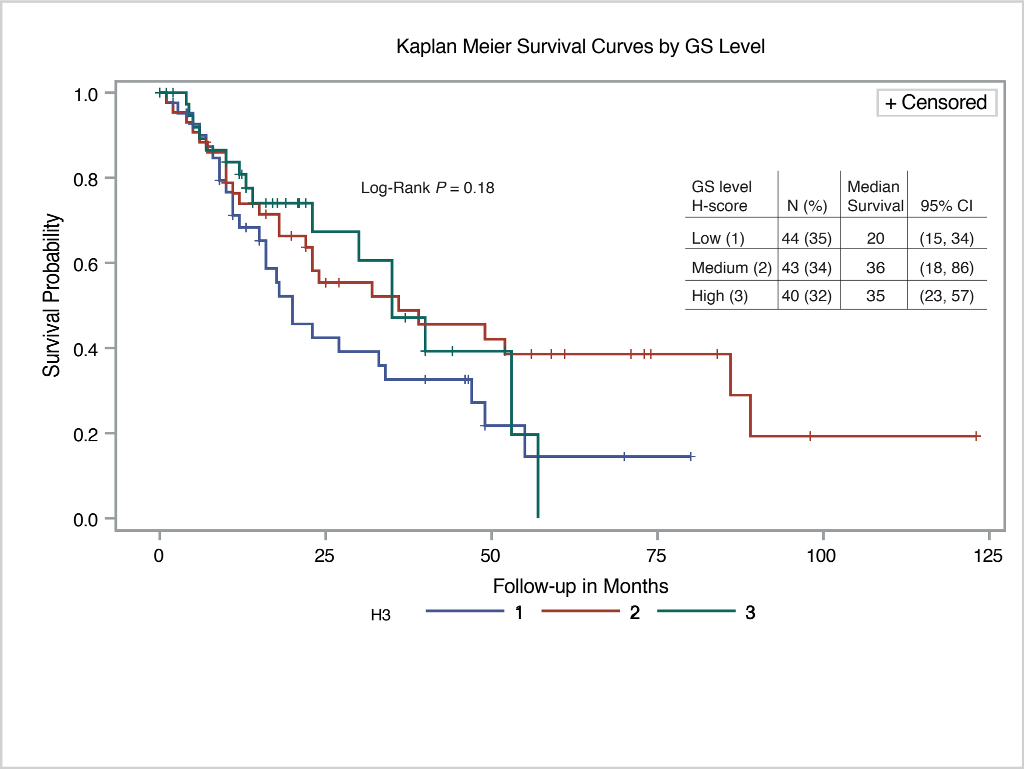
